# A fossil *Osmunda* from the Jurassic of Sweden—reconciling molecular and fossil evidence in the phylogeny of Osmundaceae

**DOI:** 10.1101/005777

**Authors:** Benjamin Bomfleur, Guido W. Grimm, Stephen McLoughlin

## Abstract

The systematic classification of Osmundaceae has long remained controversial. Recent molecular data indicate that *Osmunda* is paraphyletic, and needs to be separated into *Osmundastrum* and *Osmunda s. str*. Here we describe an exquisitely preserved Jurassic *Osmunda* rhizome (*O. pulchella* sp. nov.) that combines diagnostic features of *Osmundastrum* and *Osmunda*, calling molecular evidence for paraphyly into question. We assembled a new morphological matrix based on rhizome anatomy, and used network analyses to establish phylogenetic relationships between fossil and extant members of modern Osmundaceae. We re-analysed the original molecular data to evaluate root-placement support. Finally, we integrated morphological and molecular data-sets using the evolutionary placement algorithm. *Osmunda pulchella* and five additional, newly identified Jurassic *Osmunda* species show anatomical character suites intermediate between *Osmundastrum* and *Osmunda.* Molecular evidence for paraphyly is ambiguous: a previously unrecognized signal from spacer sequences favours an alternative root placement that would resolve *Osmunda s.l.* as monophyletic. Our evolutionary placement analysis identifies fossil species as ancestral members of modern genera and subgenera. Altogether, the seemingly conflicting evidence from morphological, anatomical, molecular, and palaeontological data can be elegantly reconciled under the assumption that *Osmunda* is indeed monophyletic; the recently proposed root-placement in Osmundaceae—based solely on molecular data—likely results from un- or misinformative out-group signals.

## INTRODUCTION

The royal ferns (Osmundales) comprise about 20 extant species currently classified in four genera, i.e. *Osmunda* L., *Osmundastrum* C.Presl, *Leptopteris* C.Presl, and *Todea* Bernh. This small group of ferns is remarkable in many respects and, consequently, has attracted considerable scholarly attention. Its members represent the most primitive of all leptosporangiate ferns (e.g. Pryer *et al.*, 2004; Smith *et al.*, 2006, 2008; Schuettpelz & Pryer, 2007), with features that have been interpreted to be intermediate between Eusporangiatae and Leptosporangiatae (e.g. Bower, 1891, 1926; Tidwell & Ash, 1994). Detailed investigations of their anatomy (e.g. Faull, 1901, 1909; Seward & Ford, 1903; Hewitson, 1962), cytology and genetic structure (e.g. Strasburger, 1900; Yamanouchi, 1910; Digby, 1919; Sharp, 1920; Manton, 1939, 1945; Manton & Smiles, 1943; Tatuno & Yoshida, 1966, 1967; Klekowski, 1970, 1973; Yatabe *et al.*, 2009), and evolution (e.g. Kidston & Gwynne-Vaughan, 1907–1910, 1914; Miller, 1967, 1971; Yatabe, Nishida & Murakami, 1999; Metzgar *et al.*, 2008; Escapa & Cúneo, 2012) render the Osmundales one of the most intensively studied groups of ferns. Moreover, in contrast to their rather limited diversity today, Osmundales have a uniquely rich and diverse fossil record (e.g. Arnold, 1964; Miller, 1971) currently considered to include more than 150 species, over 25 genera, and at least three families (e.g. Tidwell & Ash, 1994; Tian, Wang & Jiang, 2008; Wang *et al.*, 2014). This extensive fossil record has been reviewed in several key works (Miller, 1971; Tidwell & Ash, 1994; Tian *et al.*, 2008; Wang *et al.*, 2014).

The monophyly of Osmundales and their isolated position as the first diverging lineage within leptosporangiate ferns are firmly established (see, e.g. Hasebe *et al.*, 1995; Schneider *et al.*, 2004; Pryer *et al.*, 2004; Smith *et al.*, 2008). However, the resolution of systematic relationships within the group—and especially the circumscription of *Osmunda*—continues to remain controversial. Linnaeus established *Osmunda* with three species: *O. regalis* L., *O. claytoniana* L. and *O. cinnamomea* L. (Linnaeus, 1753). With subsequent descriptions of additional species from East and Southeast Asia (Thunberg, 1784; Presl, 1825; Blume, 1828; Hooker, 1837), the genus was subdivided into several subgenera, i.e. *O.* subgenus *Osmunda*, *O*. subgenus *Plenasium* (C.Presl) J.Smith, *O*. subgenus *Osmundastrum* (C.Presl) C.Presl, and *O*. subgenus *Claytosmunda* Y.Yatabe, N.Murak. & K.Iwats. based on combinations of diagnostic characters and, more recently, molecular phylogenetic analyses (Yatabe *et al.*, 1999; Yatabe, Murakami & Iwatsuki, 2005; Metzgar *et al.*, 2008). However, independent lines of evidence based on morphology (e.g. Tagawa, 1941; Hewitson, 1962; Bobrov, 1967), anatomy (Hewitson, 1962; Miller, 1967, 1971), palynology (Hanks & Fairbrothers, 1981), hybridization experiments (Tryon, 1940; Klekowski, 1971; Wagner *et al.*, 1978; Kawakami, Kondo & Kawakami, 2010), and molecular and genetic studies (e.g. Petersen & Fairbrothers, 1971; Stein & Thompson, 1975, 1978; Stein, Thomson & Belfort, 1979; Li & Haufler, 1994; Yatabe *et al.*, 1999; Metzgar *et al.*, 2008) have led to divergent opinions on the classification of these taxa; most controversy has arisen concerning the phylogenetic relationships and taxonomic ranks of *O. cinnamomea* and *O. claytoniana*.

Early molecular studies aiming to resolve specific relationships between *O. regalis*, *O. claytoniana* and *O. cinnamomea* produced remarkably incongruent results (see, e.g. Peterson & Fairbrothers, 1971; Stein & Thompson, 1975; Stein *et al.*, 1979, 1986). Isozyme studies eventually demonstrated that *O. claytoniana* is probably more closely related to *O. regalis* than either is to *O. cinnamomea* (Li & Haufler, 1994), confirming previous assumptions of early plant anatomists (e.g. Faull, 1901; Miller, 1967, 1971). Subsequent nucleotide sequencing not only provided first robust support for this relationship (Yatabe *et al.*, 1999) but, unexpectedly, also placed *Todea* and *Leptopteris* within *Osmunda* as traditionally defined. Consequently, the isolated *O. cinnamomea* at the base of the resulting tree was separated from *Osmunda s. str.* and assigned to its own genus, sister to *Leptopteris* plus *Todea* and the remaining *Osmunda* (Metzgar *et al.*, 2008; see Schuettpelz & Pryer, 2008; Smith *et al.*, 2008).

Here we describe a new *Osmunda* species based on an exceptionally well-preserved rhizome from the Jurassic of Sweden that combines diagnostic features of *Osmunda* and *Osmundastrum*. A phylogenetic analysis based on a revised morphological character matrix places the new species intermediate between *Osmunda* and *Osmundastrum*, which is incompatible with the recently established paraphyly and resulting classifications. Therefore, we re-analyse the molecular data and integrate morphological and molecular data-sets to show that the recently established paraphyly of *Osmunda s.l.* results from ambiguous outgroup signals; instead, all evidence can be elegantly reconciled assuming that *Osmunda s.l.* is indeed monophyletic.

## MATERIAL AND METHODS

### Fossil material

The studied specimen was collected from mafic volcaniclastic deposits (“Djupadal formation” of Augustsson, 2001) near Korsaröd lake (Höör municipality, central Skåne, Sweden). The host strata are interpreted to be local remnants of ash falls and lahar flows that spread from a nearby volcanic centre, similar to other occurrences of mafic volcaniclastic and epiclastic deposits associated with basaltic necks in central Skåne (Norling *et al.*, 1993; Ahlberg, Sivhed & Erlström, 2003). Palynological analyses indicate a late Pliensbachian (later Early Jurassic) age (Bomfleur, Vajda & McLoughlin, *in press*), which agrees well with radiometric dating of associated basaltic necks that place the peak phase of volcanism in central Skåne in the Pliensbachian to Toarcian (*ca* 183 Ma; Bergelin, 2009). Petrographic thin sections (Figs 1–4) were studied and photographed using an Olympus BX51 compound microscope with an attached Olympus DP71 digital camera. Two sectioned blocks of the holotype were selected for SEM analyses; the sectioned surfaces of these blocks were polished, etched with 5% HCl for 5–10 seconds, mounted on aluminium stubs, coated with gold for 90 seconds, and finally analysed using a Hitachi S-4300 field emission scanning electron microscope at the Swedish Museum of Natural History (Fig. 5). We applied conventional adjustments of brightness, contrast, and saturation to most of the digital images using Adobe^^®^^ Photoshop^^®^^ CS5 Extended version 12.0; in some cases, we performed manual image stitching and image stacking (see Kerp & Bomfleur, 2011) in order to obtain sufficiently sharp, large composite images with optimal depth of field.

**FIG. 1.**
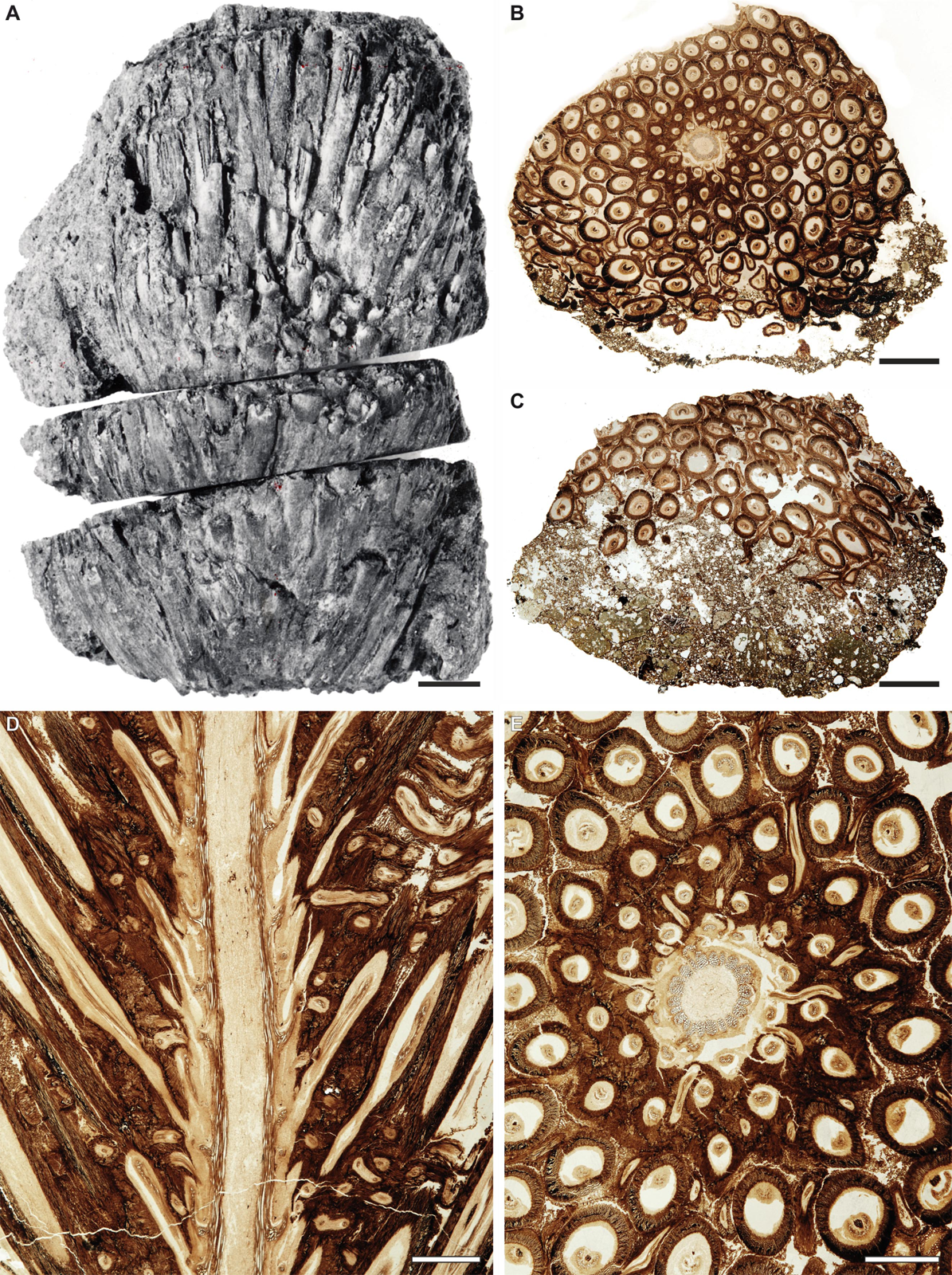
*Osmunda pulchella* sp. nov. from the Lower Jurassic of Skåne, southern Sweden. Holotype. (A) Reproduction of the only available print of the original holotype material prior to preparation, showing the gross morphology of the rhizome. (B, C) Transverse sections through center (B: NRM-S069656) and apex (C: NRM-S069657) of the rhizome. (D) Longitudinal section through the rhizome (NRM-S069658). (E) Detail of Fig. 1B. Scale bars: (A–C) = 5 mm; (D, E) = 2 mm.

**FIG. 2.**
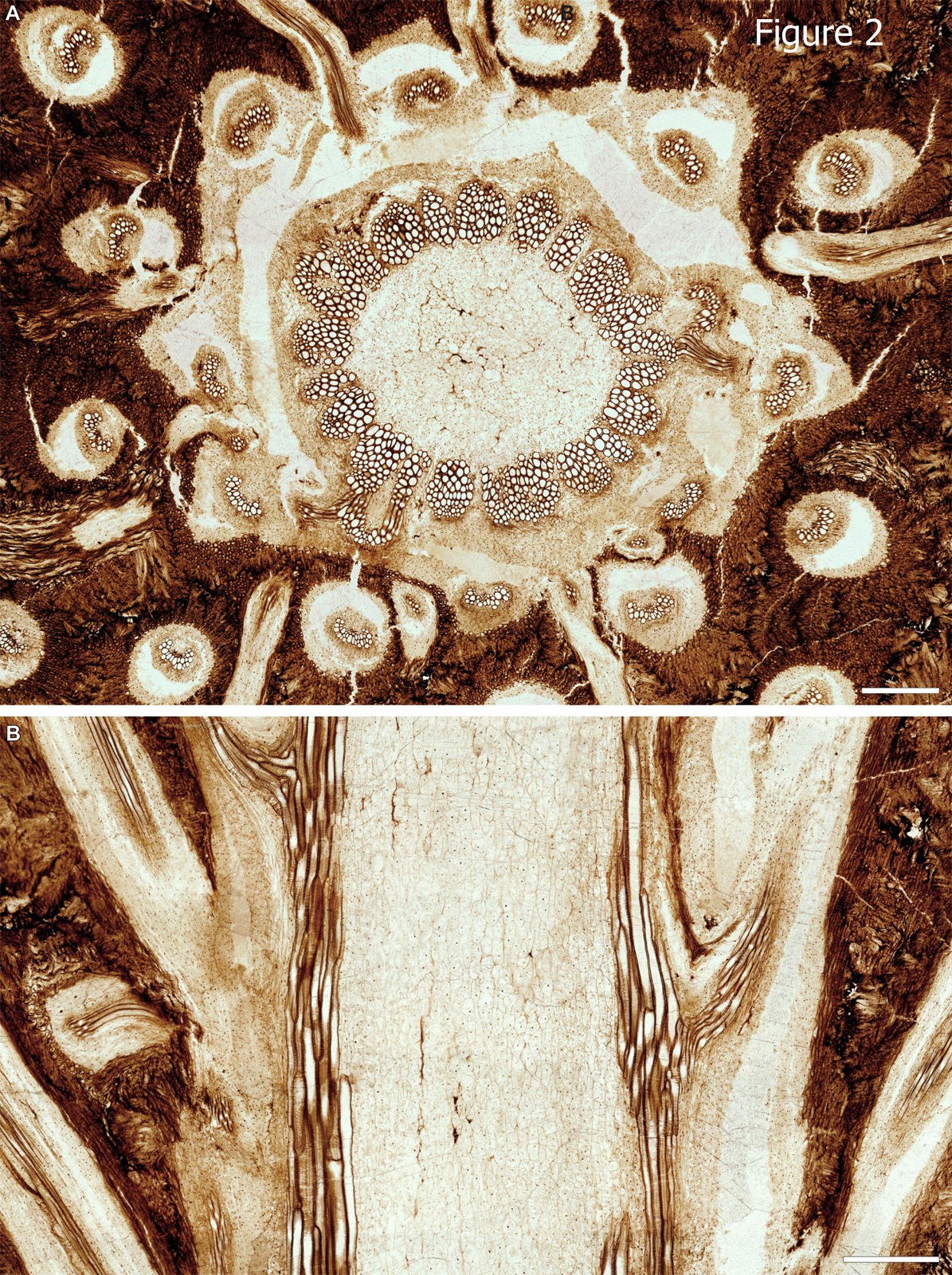
*Osmunda pulchella* sp. nov. from the Lower Jurassic of Skåne, southern Sweden. Transverse section (NRM-S069656) and radial longitudinal section (NRM-069658) through the stem. Scale bars = 500 µm.

**FIG. 3.**
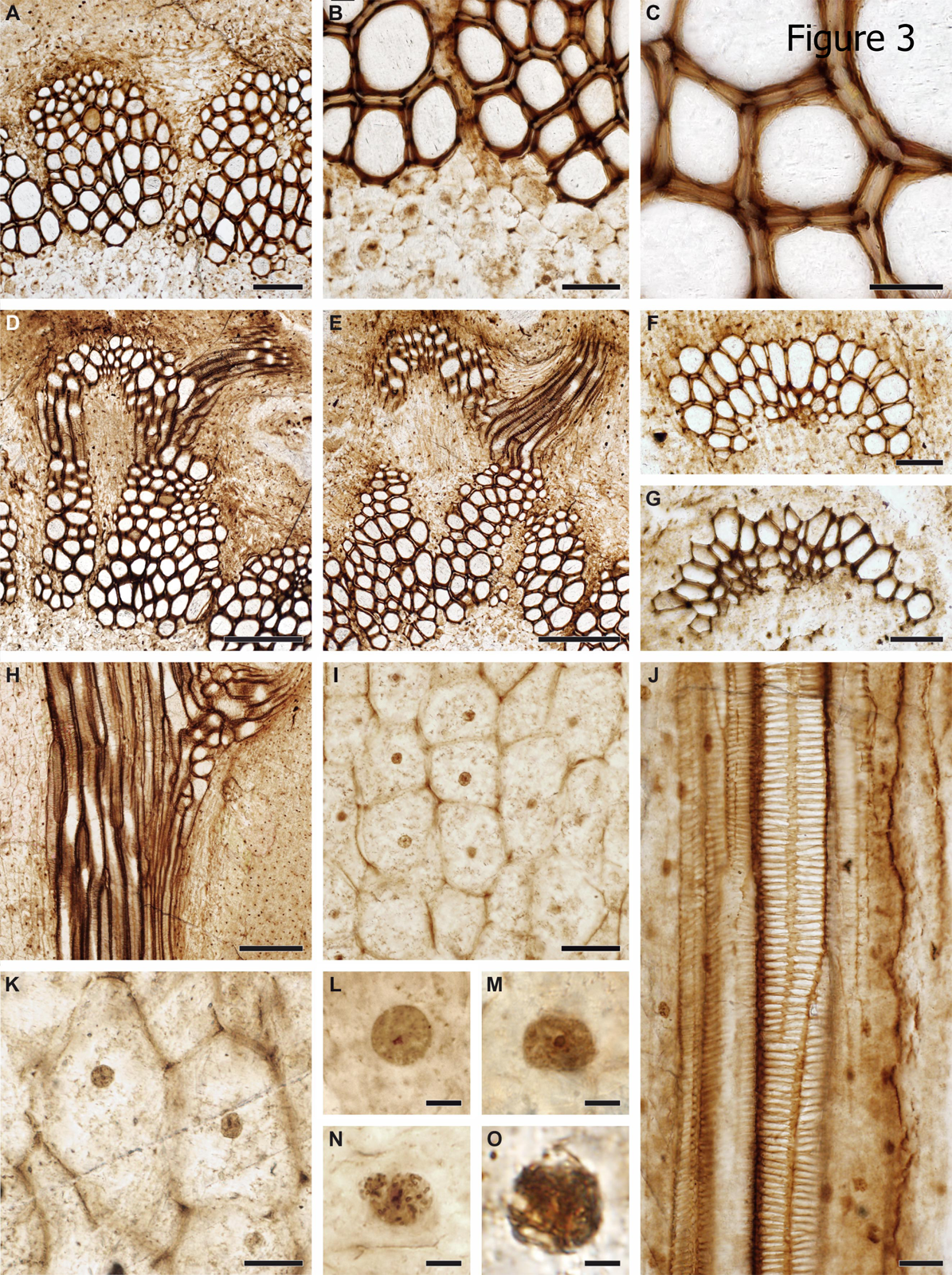
Anatomical and cytological details of *Osmunda pulchella* sp. nov. from the Lower Jurassic of Skåne, southern Sweden (A–G, L–N: Cross-section, NRM-S069656; H–K, O: Radial longitudinal section, NRM-S069658). (A) Detail showing pith parenchyma (bottom), stelar xylem cylinder dissected by complete leaf gaps, triangular section of phloem projecting into leaf gap, and parenchymatous inner cortex (top); note mesarch leaf-trace protoxylem initiation in the stelar xylem segment on the right. (B) Detail of (A) showing peripheral pith parenchyma and stem xylem. (C) Detail of stem xylem showing tracheid pitting. (D, E) Endarch leaf traces emerging from the stele, each associated with a single root. (F) Leaf trace in the inner cortex of the stem showing single, endarch protoxylem cluster. (G) Leaf trace immediately distal to initial protoxylem bifurcation in the outermost cortex of the stem. (H) Detail showing pith parenchyma (left), stelar xylem cylinder with emerging leaf trace (centre), and parenchymatous inner cortex (right). (I, K). Well-preserved pith parenchyma showing membrane-bound cytoplasm with cytosol particles and interphase nuclei containing nucleoli. (J) Root vascular bundle with well-preserved scalariform pitting of metaxylem tracheid. (L, M, N, O) Nuclei with conspicuous nucleoli (in interphase: L, M) or with condensed chromatin or distinct chromatid strands (in prophase: N, O). Scale bars: (A) = 100 µm; (B, F, G, I) = 50 µm; (C, J, K) = 25 µm; (D, E, H) = 200 µm; (L–N) = 5 µm; (O) = 2.5 µm.

**FIG. 4.**
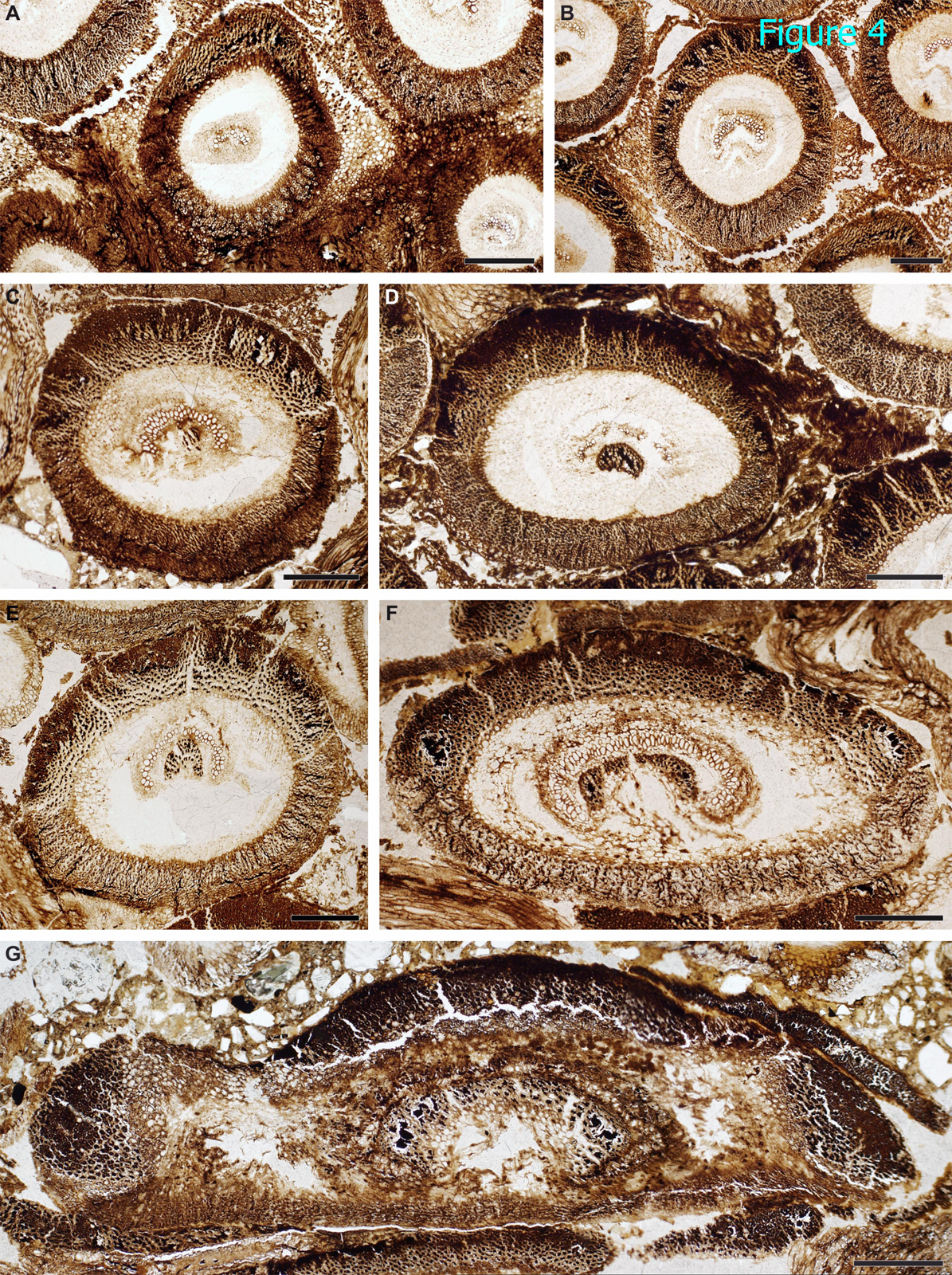
Basal to distal sections of petiole bases of *Osmunda pulchella* sp. nov. from the Lower Jurassic of Skåne, southern Sweden (NRM-S069657), showing successive stages of petiole-base differentiation. (A) Inner cortex and petiolar wings parenchymatous. (B–E) Development of an abaxial arch of thick-walled fibres in the sclerenchyma ring. (C) Appearance of a sclerenchyma patch in the bundle concavity. (D) Appearance of a sclerenchymatic mass in the petiolar wing. (F) Sclerenchyma ring with two prominent lateral masses of particularly thick-walled fibres. (G) Collapsed outermost petiole (note rock matrix above) showing sclerenchyma ring with one abaxial and two lateral masses of particularly thick-walled fibres, and elongate sclerenchyma strips (e.g. bottom, right) isolated from degraded stipular wings of adjacent petioles. Scale bars = 500 µm.

**FIG. 5.**
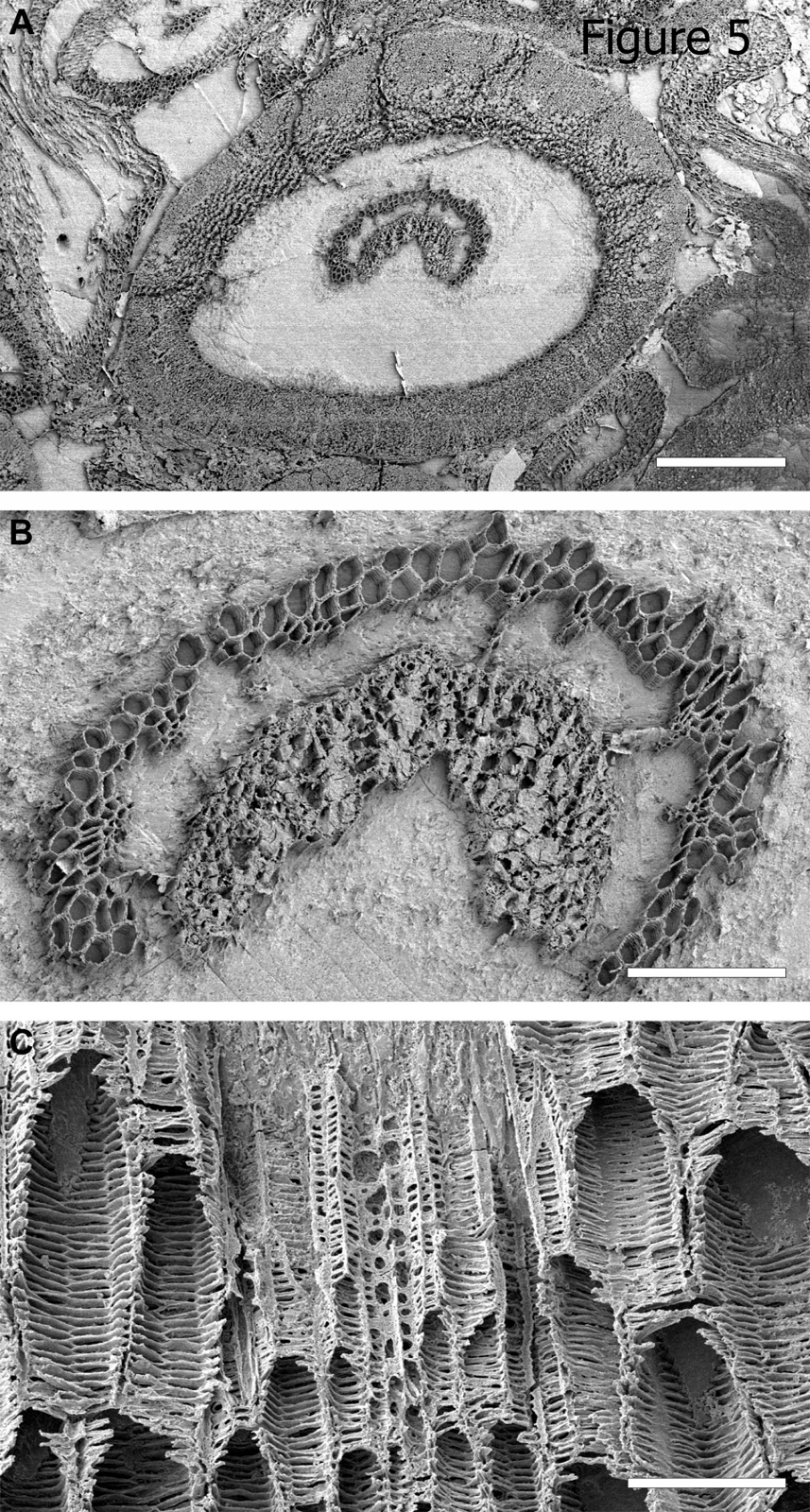
Details of petiole-base anatomy of *Osmunda pulchella* sp. nov. from the Lower Jurassic of Skåne, southern Sweden, revealed via scanning electron microscopy. (A) Distal cross-section through a petiole. (B) Detail of (A) showing vascular strand with about eight endarch protoxylem bundles and sclerenchyma mass lining the vascular-strand concavity. (C). Detail showing helical wall thickening of protoxylem strands (center) compared to multiseriate scalariform wall thickenings of metaxylem tracheids in a petiole vascular bundle (oriented with adaxial side facing upwards). Scale bars: (A) = 1 mm; (B) = 100 µm; (C) = 50 µm.

### Phylogenetic analyses

In order to place the newly described fossil in a phylogenetic context, we assembled a morphological matrix that is based on the phylogenetic assessment of Miller (1971), including all extant and fossil members of the extant genera (Wang *et al.*, 2014) **[Supplementary Information]**.

*Network analysis. —* We rely exclusively on network methods as implemented in SplitsTree v. 4.13.1 (Huson & Bryant, 2006) to draw phylogenetic conclusions based on the morphological matrix (see Spencer *et al.,* 2004; Denk & Grimm, 2009; Friis *et al.*, 2009; Schlee *et al.*, 2011; Grímsson *et al.,* 2014): (1) a neighbour-net (Bryant & Moulton, 2002, 2004) based on mean inter-taxon distances, and (2) bipartition networks to visualize support (Bayesian-inferred posterior probabilities, PP; non-parametric bootstrapping, BS) for alternative phylogenetic relationships (Holland & Moulton, 2003; Grimm *et al.*, 2006; Denk & Grimm, 2009). BS support was established under three commonly used optimality criteria using 10,000 bootstrap replicates: (1) Least-squares via the BioNJ algorithm (BS_NJ_; Gascuel, 1997); (2) Maximum parsimony (BS_MP_) using PAUP* (Swofford, 2002; Müller, 2005); and (3) Maximum likelihood (BS_ML_) via the fast bootstrapping implementation in RAxML v. 7.4.2 (Stamatakis, 2006; Stamatakis, Hoover & Rougemont, 2008) using both available transition models for categorical (multistate) data, i.e. (*i*) the general time-reversible model (BS_ML/GTR_) (Rodriguez *et al.*, 1990) and (*ii*) Lewis’ (2001) model (BS_ML/MK_). For set-up details of Bayesian inference, non-parametric bootstrapping, and network-wise visualisation refer to File S1 **[Supplementary Information]**.

*Re-visiting the Osmundaceae root. —* We analysed the root placement in the phylogenetic tree of Metzgar *et al.* (2008) using the original molecular matrix. First, a set of traditional phylogenetic analyses was run, including a gene jackknifing procedure. Trees and bootstrap support were inferred using the concatenated data, each gene partition separately, and matrices in which one partition was deleted. Second, the evolutionary placement algorithm (EPA; Berger & Stamatakis, 2010; Berger, Krompass & Stamatakis, 2011) as implemented in RAxML was used to determine the optimal position of the outgroup taxa (i. e. the position of an outgroup-inferred root) within an ingroup-only topology. The EPA has been originally designed for placing fossils (Berger & Stamatakis, 2010) or short-sequence reads (Berger *et al.,* 2011), but its metrics can also be used to generally test the position of one or many query sequences —here: outgroup taxa— in a given topology —here: an ingroup-only ML tree— in a ML framework (A. Stamatakis, pers. comm., 2014).

*Character plotting and independent optimisation of the placement of fossils within a molecular framework of modern taxa. —* Using the EPA we estimated a weight (probability) for the placement of our fossil within the molecular backbone topology reproduced from the data matrix of Metzgar *et al.* (2008). We also determined the most parsimonious placement of the newly described fossil within the molecular tree of Metzgar *et al.* (2008) implemented into the morphological matrix using MESQUITE v. 2.75 (Maddison & Maddison, 2011); this is done by simply moving the fossil within the given topology and recording the incremental increase in steps added to the resulting whole tree-length [see Friis *et al.* (2009) and von Balthazar *et al.* (2012) for applications]. Nomenclatural remark

In order to maintain consistent use of terminology, we employ the following names: (1) *Osmunda cinnamomea* instead of the currently used *Osmundastrum cinnamomeum* (L.) C.Presl; (2) *Osmunda chengii* nom. nov., a replacement name for a new combination that is based on *Ashicaulis claytoniites* Y.M.Cheng (see Discussion and Appendix); (3) ‘modern Osmundaceae’, referring to those genera of Osmundaceae that are based on extant species, i.e. *Osmunda* (including *Osmundastrum*), *Todea*, and *Leptopteris*; (4) ‘*Osmunda s.l.*’, referring to the traditional generic concept that includes all extant and several fossil species (e.g., Miller, 1971); and (5) ‘*Osmunda s. str.*’, referring to the recently proposed generic concept of *Osmunda* that excludes *O. cinnamomea* and *O. precinnamomea* C.N.Mill. (i.e., including only *Osmunda* subgenera *Osmunda, Claytosmunda* and *Plenasium*) (Yatabe *et al.*, 1999). Where necessary, we cite taxon authorities to discriminate between formal subgeneric concepts used by Miller (1971) and Yatabe *et al.* (2005).

## RESULTS

### Systematic description of the fossil

Order Osmundales Link

Family Osmundaceae Berch. & C.Presl

Genus *Osmunda* L.

Species Osmunda pulchella sp. nov.

*Diagnosis*. — **Rhizome** creeping or semi-erect. **Stem** with ectophloic-dictyoxylic siphonostele and two-layered cortex. **Pith** entirely parenchymatous. **Xylem cylinder** about 8–12 tracheids (mostly *ca* 0.4 mm) thick, dissected by narrow, complete, immediate leaf gaps, containing about twenty xylem segments in a given transverse section. **Phloem and endodermis** external only. **Inner cortex** *ca* 0.5–0.8 mm thick, homogeneous, parenchymatous, containing about ten leaf traces in a given transverse section; **outer cortex** *ca* 1.5–2.5 mm thick, homogeneous, sclerenchymatous, containing about 20 leaf traces in a given transverse section. **Leaf traces** in stem oblong, more or less reniform, adaxially concave, endarch with a single protoxylem strand at the point of emergence from stele, diverging at acute angles of *ca* 20–40°; protoxylem strand bifurcating only in outermost cortex or upon departure from stem. **Petiole bases** with adaxially concave vascular strand, one adaxial sclerenchyma band in vascular-strand concavity, parenchymatic cortex, a heterogeneous sclerenchyma ring, and an opposite pair of petiolar wings; **adaxial sclerenchyma** in inner cortex of petiole appearing in form of a single patch or arch lining the vascular-bundle concavity with homogeneous thickness, differentiating distally into two thickened lateral masses connected by a thin strip, extending proximally only to base of petiole, not into stem; **sclerenchyma ring** of petiole base thicker than vascular bundle, heterogeneous, with a crescentic abaxial cap of thicker-walled fibres in the basal petiole portion differentiating distally into two lateral masses and ultimately into two lateral and one abaxial mass; **petiolar wings** in distal portions containing an elongate strip of thick-walled fibres. **Roots** diarch, usually arising singly from one leaf trace, containing scattered sclerenchyma fibres.

*Type stratum and age*. — Mafic pyroclastic and epiclastic deposits informally named the “Djupadal formation”; Pliensbachian (later Early Jurassic).

*Type locality*. — Korsaröd lake (55°58′54.6”N, 013°37′44.9”E) near Höör, central Skåne, southern Sweden.

*Holotype* (*hic designatus*). — A single specimen of permineralized rhizome, sectioned and prepared into six blocks (specimens NRM S069649–S069655) and three microscope slides, including two transverse thin sections (slides NRM S069656 and S069657) and one radial thin section (NRM S069658); all material is curated in the Collection of the Department of Palaeobiology, Swedish Museum of Natural History, Stockholm, Sweden.

*Etymology*. — The specific epithet *pulchella* (lat. ‘beautiful little’) is chosen in reference to the exquisite preservation and aesthetic appeal of the holotype specimen.

*Description*. — The holotype specimen is a calcified rhizome fragment about 6 cm long and up to 4 cm in diameter (Fig. 1A–C). It consists of a small central stem that is surrounded by a compact mantle of helically arranged, persistent petiole bases and interspersed rootlets (Fig. 1B, E). The rootlets extend outwards through the mantle in a sinuous course almost perpendicular to the axis, indicating low rhizomatous rather than arborescent growth; the asymmetrical distribution of roots in longitudinal sections of the rhizome (Fig. 1D) points to a creeping habit.

The stem measures about 7.5 mm in diameter, and consists of an ectophloic-dictyoxylic siphonostele surrounded by a two-layered cortex (Figs 1D, E, 2, 6A). The pith is *ca* 1.5 mm in diameter and entirely parenchymatous (Fig. 2). A thin region at the outermost periphery of the pith consists of a few rows of parenchyma cells that are considerably more slender (*ca* 20–30 µm wide) than those in the central portion of the pith (usually ≥ 50 µm wide) (Figs 2B, 3A, B, H); furthermore, cell walls in some regions of the pith periphery may be thicker and more clearly visible than in the centre (Figs 2A, 3B). However, there is no evidence for the presence of an internal endodermis or internal phloem. Given that endodermal layers are recognisable in the stem and petiole cortices (e.g. Fig. 4F), we are positive that the absence of an internal endodermis is an original feature, and not the result of inadequate preservation. The xylem cylinder is *ca* 0.4 mm and *ca* 8–12 tracheids thick, and dissected by narrow, mostly complete, immediate leaf gaps into about 20 xylem segments in a given transverse section. The phloem forms an entire ring around the stele; it is most easily recognisable opposite a leaf gap, where it forms a narrow wedge-shaped patch of large, thin-walled cells that projects slightly towards the gap in transverse section (Figs 2A, 3A).

**FIG. 6.**
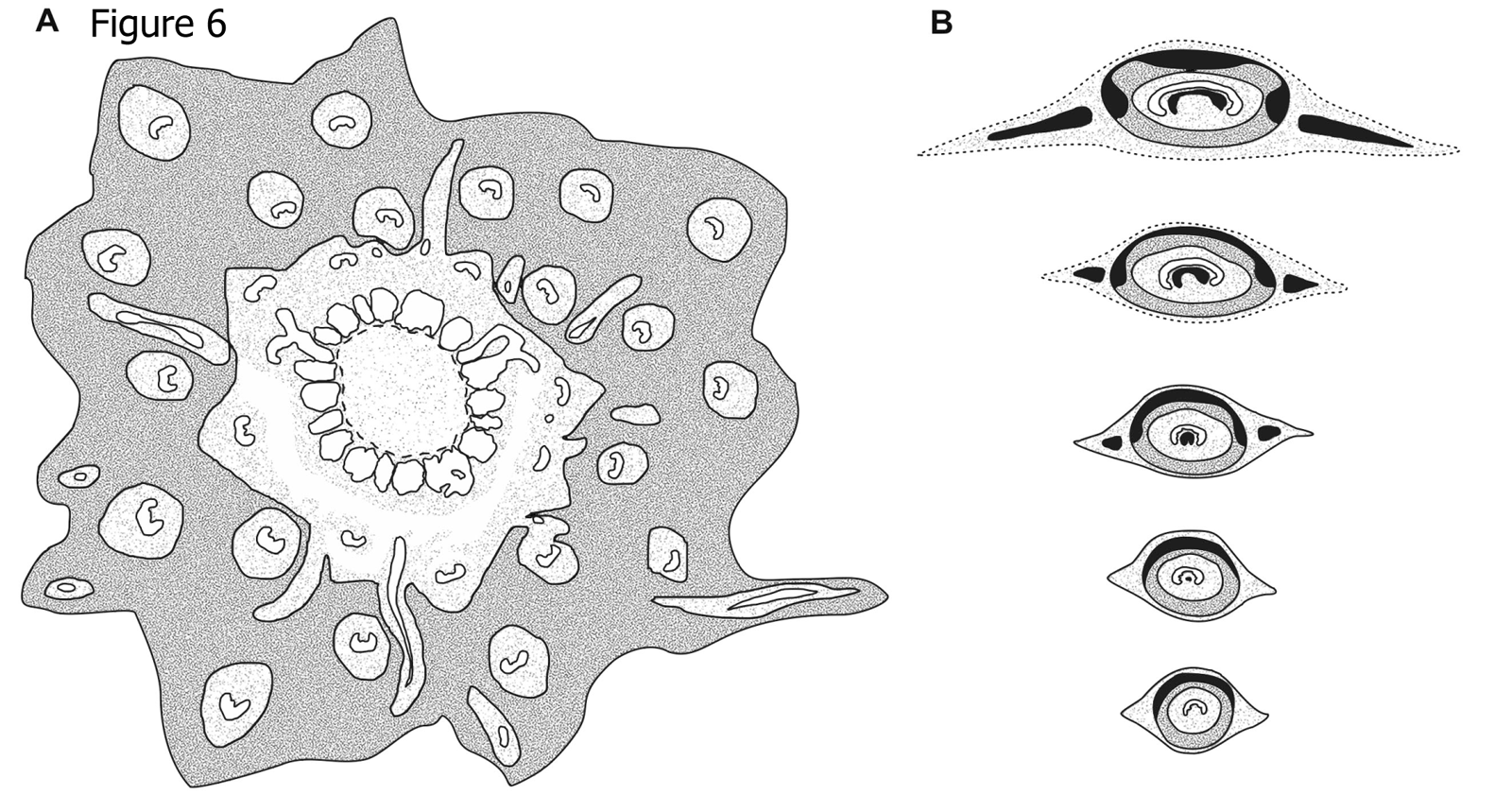
Schematic drawings showing diagnostic anatomical characters of *Osmunda pulchella* sp. nov. from the Lower Jurassic of Skåne, southern Sweden. (A) Stem cross section. (B) Successive cross sections of basal (bottom) to distal (top) petiole portions. Xylem in white; parenchyma in light-grey; sclerenchyma in dark-grey; sclerenchyma with particularly thick-walled fibres in black.

The cortex of the stem is bi-layered (Figs 1E, 2, 6A). The inner layer is *ca* 0.5–0.8 mm thick, consists entirely of parenchyma, and contains about ten leaf traces in a given transverse section (Fig. 2A). The outer cortex is considerably thicker (*ca* 1.5–2.5 mm thick), and consists entirely of homogeneous sclerenchymatic tissue (Figs 1E, 2). Abundant leaf traces (about 20 in a given transverse section; e.g. Fig. 2A) and rootlets traversing the outer cortex (Figs 1C, D, 2) appear to have altered the original orientation of the sclereids, resulting in a somewhat patchy appearance of the outer cortical tissue (Fig. 2).

Phyllotaxy of the stem is helical with apparent contact parastichies of 8 and 13 (Fig. 1B, E). Leaf-trace formation begins with the appearance of a single protoxylem strand in an eccentric position (about two-thirds to three-quarters distance from the pith; Fig. 3A) in a stelar metaxylem segment. Distally, the protoxylem becomes associated with an increasing amount of parenchyma on its adaxial side (making it effectively endarch for the rest of its course), first occupying only the centre of the segment (resulting in an O-shaped xylem segment), then connecting with the pith (resulting in a U-shaped xylem segment), and ultimately forming the usually complete, narrow leaf gap with the departure of the trace. Departing leaf traces are oblong, only slightly curved adaxially, *ca* 300–350 µm wide and two to four tracheids (*ca* 80–100 µm) thick (Figs 2, 3D, E), and diverge from the axis at angles of *ca* 20–40° (Figs 1D, 2B).

In its course through the stem, a leaf-trace vascular bundle becomes enveloped by increasing layers of tissue through which it successively passes: first by phloem and endodermis from the stele upon entering the inner cortex; by a sheath of parenchyma from the inner cortex as it enters the outer cortex (Fig. 2); and finally by a cylindrical sclerenchyma sheath from the outer cortex as it departs from the stem (Fig. 1E). The initial bifurcation of the leaf-trace protoxylem occurs in the outermost portion of the cortex or in the petiole base (Fig. 3F, G).

In the inner cortex of the petiole, thick-walled fibres appear in form of a small irregular mass adaxial to the vascular bundle (Fig. 4C, D). This develops distally into a thick band lining the bundle concavity (Figs 4E, 5A, B), and may further differentiate into two lateral masses connected only by a rather thin strip (Fig. 4F, G). Apart from the sclerenchyma inside the vascular-bundle concavity, the inner cortex of the petiole consists entirely of parenchyma.

The sclerenchyma cylinder of the petiole has an even thickness that increases from about 300 µm near the petiole base to *ca* 500 µm distally. Its composition is heterogeneous: near the petiole base, it contains a crescentic, abaxial arch of particularly thick-walled fibres (Figs 1E, 4, 5, 6B); distally, this arch begins to develop two lateral masses (Figs 4D–F, 6B) and ultimately two lateral masses and one abaxial arch of thick-walled fibres whose lumina are more-or-less entirely occluded (Figs 4G, 6B).

The petiole bases are flanked by a pair of stipular wings that consist initially of parenchyma only; as the wings grow wider in more distal portions, they develop a patch of thick-walled fibres (Figs 4D, 6B) that forms an entire, elongate strip (Figs 4F, G, 6B). The parenchymatic ground tissue of the stipular wings is well-preserved only in the innermost regions of the mantle (Figs 1B, C, E); outwards, it appears to be either increasingly degraded or to have been removed by the abundant penetrating rootlets. In the outermost portions of the mantle, all that remains of the stipular wings are usually just the isolated, elongate strips of thick-walled fibres interspersed between petioles and rootlets (Fig. 4G).

Each leaf trace is usually associated with a single rootlet that diverges laterally at the point of departure from the stele. The rootlets typically measure about 0.5 mm in diameter, contain a diarch vascular bundle, parenchymatic ground tissue with interspersed sclerenchymatic fibres, and a sclerenchymatic outer cortical layer.

The holotype specimen of *O. pulchella* shows a phenomenal quality of preservation of cellular and subcellular detail (Figs 2, 3). Tracheids have exquisitely preserved wall thickenings, which are scalariform in metaxylem (Figs 3C, J, 5C) and annular to helical in protoxylem cells (Fig. 5C). Most parenchyma cells contain preserved cellular contents (Figs 2, 3), including nuclei (Fig. 3H–K), membrane-bound cytoplasm (Fig. 3I), and cytosol granules (Fig. 3I, K) (see Bomfleur *et al.*, *in press*). Some parenchyma cells, especially those adjacent to xylem bundles in roots and leaf traces, contain varying amounts of discrete, smooth-walled, spherical or oblate particles *ca* 1–5 µm in diameter that have been interpreted as putative amyloplasts (Bomfleur *et al.*, *in press*). Cell nuclei measure *ca* 10 µm in diameter, and contain nucleoli and chromosomes (Fig. 3K–O). Chromatid strands have a diameter of 0.3–0.4 µm (Fig. 3O).

### Phylogenetic analyses

*Phylogenetic relationships among fossil and modern members of the Osmundaceae based on rhizome anatomy (Fig. 7)*. — The phylogenetic network based on pairwise distances inferred from a matrix including 23 rhizome anatomical characters resolved five major species groups: (1) extant species of *Leptopteris* and *Todea* together with *T. tidwellii* Jud, G.W.Rothwell & Stockey from the Lower Cretaceous of North America; (2) all extant species of *Osmunda* subgenus *Plenasium* together with *O. arnoldii* C.N.Mill. and *O. dowkeri* (Carruth.) M.Chandler from the Paleogene of North America and Europe; (3) all species of subgenus *Osmunda sensu* Miller, i.e. species of the extant subgenera *Osmunda sensu* Yatabe *et al.* and *Claytosmunda* together with several Paleogene and Neogene species; (4) the Jurassic *Osmunda* species, including *O. pulchella*; and (5) all extant and fossil members of *Osmundastrum* (incl. *O. precinnamomea*). Corresponding bipartitions, which would define clades in an accordingly rooted phylogram, were found in the bootstrap replicate tree sample and the Bayesian sampled topologies with varying frequency. *Osmunda* subgenus *Plenasium* (BS = 47–80; PP = 0.76) and *Osmundastrum* (BS = 51–76; PP = 0.95) received best support, whereas support values for the other groups were generally low (BS ≤ 55; PP ≤ 0.43). The Jurassic species bridge the morphological gap between *Osmundastrum* and *O.* subgenus *Osmunda sensu* Miller, with *O. pulchella* being the species closest to *Osmundastrum*. A hypothetical clade comprising *O.* subgenus *Osmunda sensu* Miller and the Jurassic *Osmunda* species would receive BS up to 28 and PP of 0.28.

**FIG. 7.**
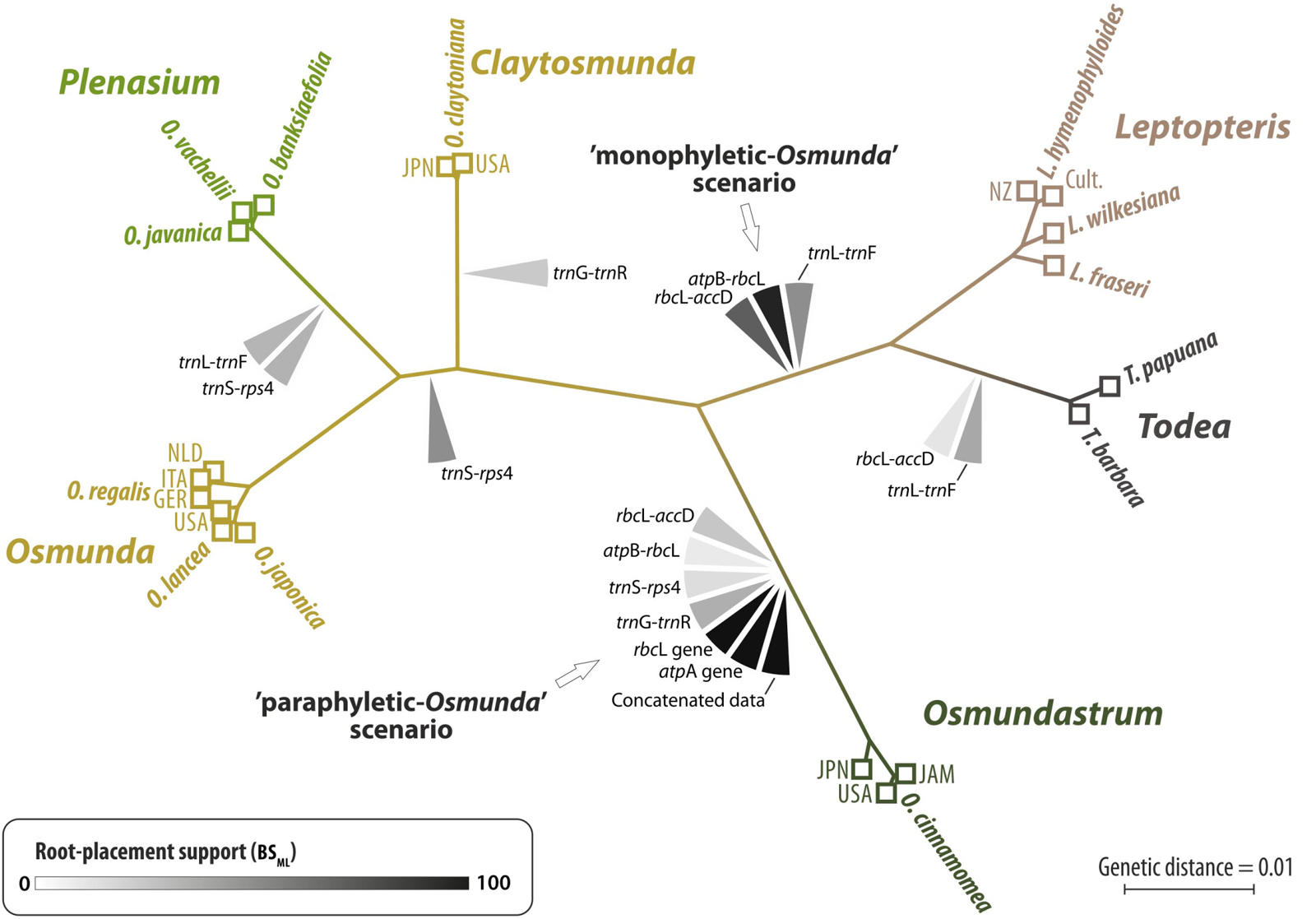
Neighbour-net showing phylogenetic relationships among fossil and extant members of modern Osmundaceae inferred from a morphological distance matrix based on rhizome anatomy. Edge (branch) support from bootstrapping (BS) and Bayesian inference (posterior probability, PP) is annotated for modern genera and subgenera, and selected bipartitions. Further abbreviations: BS_ML/GTR_, maximum likelihood (ML) BS support, using a general-time reversible transformation model; BS_ML/MK_, BS support, using Lewis’ (2001) one-parameter model; BS_P_, parsimony BS support; BS_NJ_, neighbour-joining BS support. **[Supplementary Information]**.

Especially remarkable is the diversification of subgenus *Osmundastrum* as revealed by our independent coding of the individual fossil records from Neogene (Miller, 1967; Matsumoto & Nishida, 2003), Paleogene (Miller, 1967), and Cretaceous deposits (Serbet & Rothwell, 1999); the individually coded fossil and extant representatives assigned to *O. cinnamomea* show greater morphological disparity than expressed between the separate species of any other subgenus and genus.

*Merging fossil and extant taxa into a molecular backbone topology (Fig. 8). —* Of all taxa placed via EPA (evolutionary placement algorithm), *Osmunda pulchella* is the species that is most incongruently placed between the different weighting schemes: Using parsimony-based character weights, the EPA places *Osmunda pulchella* at the root of *Claytosmunda*, whereas it is placed either between *Osmundastrum* and the remaining *Osmunda s. str.* or at the root of the *Plenasium* clade using model-based character weights. Single position swaps also occur in most of the other Jurassic species [*O. plumites* (N.Tian & Y.D.Wang) comb. nov., *O. wangii* (N.Tian & Y.D.Wang) comb. nov., *O. johnstonii* (Tidwell, Munzing & M.R.Banks) comb. nov., *O. liaoningensis* (Wu Zhang & Shao-Lin Zheng) comb. nov.] and in *O. pluma* C.N.Mill., *O. iliaensis* C.N.Mill., *O. shimokawaensis* M.Matsumoto & H.Nishida, and *Todea tidwellii*. Except for *Todea tidwellii* (placed at the root of either *Leptopteris* or *Todea*), all swaps occur within the *Osmunda s.l.* sub-tree. Swaps among the Jurassic species mostly involve placements at the root of the *Plenasium* sub-tree, the subgenus *Osmunda* sub-tree, and at the branch between *Osmundastrum* and the remaining *Osmunda*. *Osmunda shimokawaensis* and *O. iliaensis* are variably placed within the *O. lancea* Thunb.*–O. japonica* Thunb. sub-tree.

**FIG. 8.**
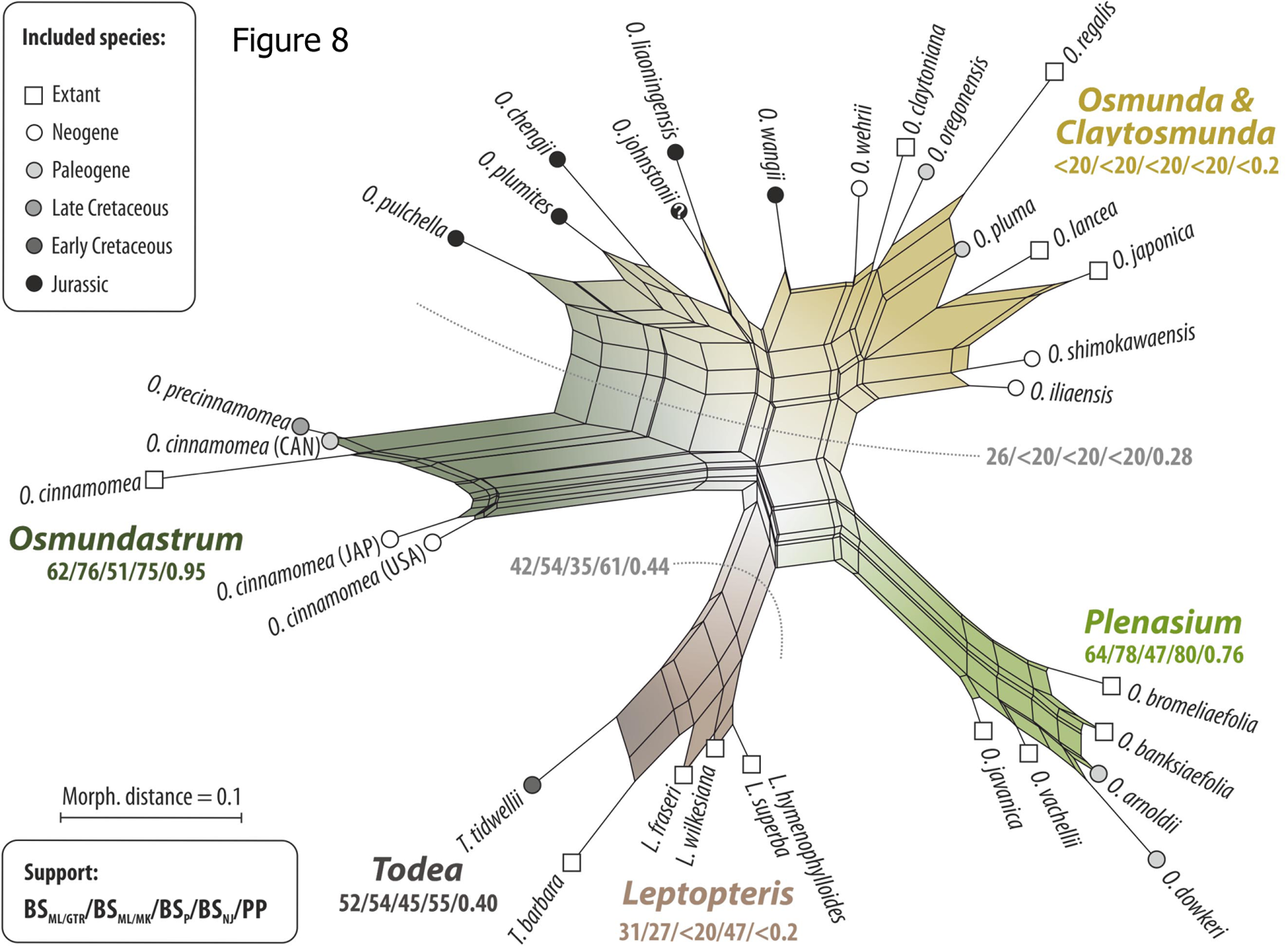
Placement of fossil and extant members into the specified backbone topology of modern Osmundaceae inferred from molecular data of Metzgar *et al.* (2008) using the evolutionary placement algorithm (Berger and Stamatakis, 2010) **[Supplementary Information]** and three different character-weighting schemes; dashed light-grey lines indicate weighting-scheme-dependent position swaps of taxa. Abbreviations: ML_GTR_, weighting scheme for morphological characters optimized under a general-time reversible transformation model; ML_MK_, weighting scheme optimized under

By contrast, fixed placements congruent over all three weighting schemes employed occur in: Fossil members of *Osmundastrum* (all at the *O. cinnamomea* branch); *O. chengii* and *O. wehrii* C.N.Mill. (at the root of the *Plenasium* sub-tree); *O. arnoldii*, *O. bromeliaefolia* (C.Presl) Copel., and *O. dowkeri* (all at *O. banksiaefolia* branch); *O. oregonensis* (C.A.Arnold) C.N.Mill. (at the root of subgenus *Osmunda*), and *L. superba* (at the branch of *L. hymenophylloides*).

*Re-visitation of the outgroup-inferred Osmundaceae root (Fig. 9)*. — The gene jackknifing and single-gene analyses reveal ambiguity concerning the position of the Osmundaceae root in the data of Metzgar *et al.* (2008). As in the original analysis, support for backbone branches is effectively unambiguous based on the concatenated data, and places the outgroup between *Osmundastrum* and the remainder of the family, resolving the traditional genus *Osmunda* (*Osmunda s.l.*) as a grade (‘paraphyletic *Osmunda* scenario’). The signal for this root placement stems from the two coding plastid gene regions (*atp*A and *rbc*L). In the more (but not most) variable spacer regions (*rbc*L-*acc*D, *atp*B-*rbc*L, and *trn*L-*trn*F to a lesser degree), however, a competing signal is found resolving genus *Osmunda s.l.* as a clade (‘monophyletic *Osmunda* scenario’). The most variable non-coding spacer regions (*trn*G-*trn*R; *rps*4-*trn*S; and *trn*L-*trn*F to some degree) provided only ambiguous signals including potential outgroup-branch placements deep within the *Leptopteris-Todea* and *Osmunda* sub-trees and showed a preference for an *Osmundastrum-Leptopteris-Todea* clade as sister to *Osmunda s. str*.

**FIG. 9.**
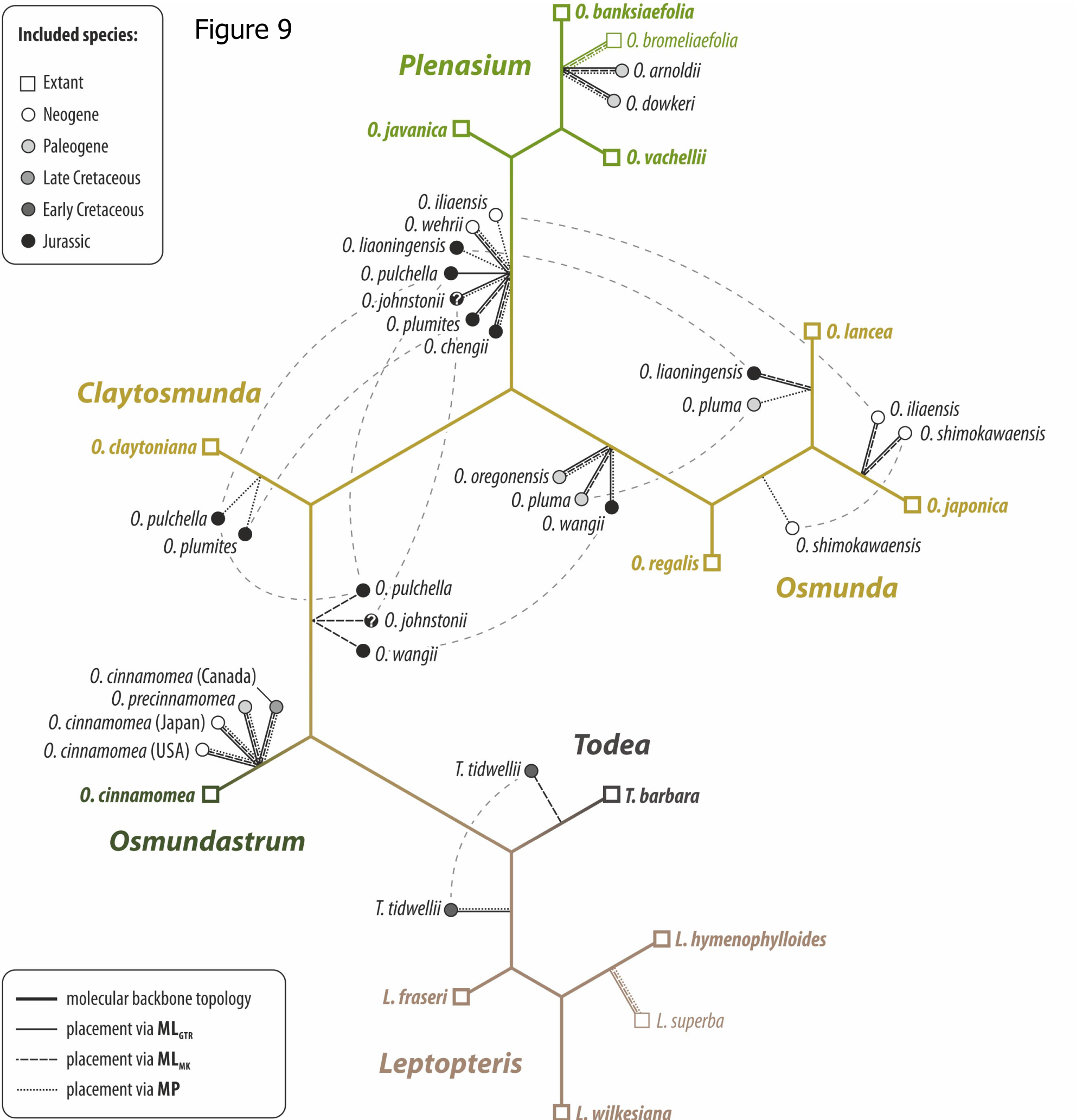
Phylogenetic tree, optimised under maximum likelihood (ML), showing unambiguously resolved relationships among extant Osmundaceae and the conflicting root-placement (outgroup-inferred) signals from individual gene regions; based on the molecular matrix compiled and employed by Metzgar *et al.* (2008). All backbone branches received full maximum-likelihood bootstrap support (BS_ML_ = 100) based on the concatenated data; support for leaf-branches not shown (see **[Supplementary Information]**).

The gene-jackknifing results showed that the exclusion of either one or both coding regions (*atp*A, *rbc*L)—which together account for 33% of distinct alignment patterns in the concatenated matrix—decreased support for the split leading to an *Osmunda* grade with *Osmundastrum* resolved as sister to the remainder of the family, whereas the support for the alternative of an *Osmunda* clade or an *Osmundastrum-Leptopteris-Todea* clade was increased. In the case of *O. (Claytosmunda) claytoniana*, the genetic data provided a coherent signal, with all plastid regions preferring a subgenus *Osmunda sensu* Yatabe *et al.–Plenasium* clade over the alternatives of a subgenus *Osmunda sensu* Miller or *Claytosmunda-Plenasium* clade.

The gene-knifing had no measurable effect (BS_ML_ = 98–100). The problem with the placement of the root can also be illustrated in form of a neighbour-net splits graph based on genetic, uncorrected p-distances [see Fig. S1 in Electronic Supplementary Archive (ESA)].

*Placement of* Osmunda pulchella *within the two molecular backbone topologies (Fig. 10). —* Optimisation of the anatomical characters on two specified backbone topologies inferred from the different rooting scenarios (‘monophyletic *Osmunda*’ *vs* ‘paraphyletic *Osmunda*’ scenario) required 53 steps under parsimony. Inserting *Osmunda pulchella* into the ‘paraphyletic *Osmunda* scenario’ tree, its most parsimonious placement based on anatomical characters is either (1) at the most basal position as sister to all extant Osmundaceae, (2) as sister to *O. cinnamomea*, or (3) as sister to a putative *Leptopteris-Todea-Osmunda s. str.* clade. In the ‘monophyletic *Osmunda* scenario’ tree, by contrast, the most parsimonious placement of *O. pulchella* is as sister to *O. cinnamomea*. In both trees, the least parsimonious positions of *O. pulchella* are within the *Todea*-*Leptopteris* clade or at the root of or within the *Plenasium* subtree.

**FIG. 10.**
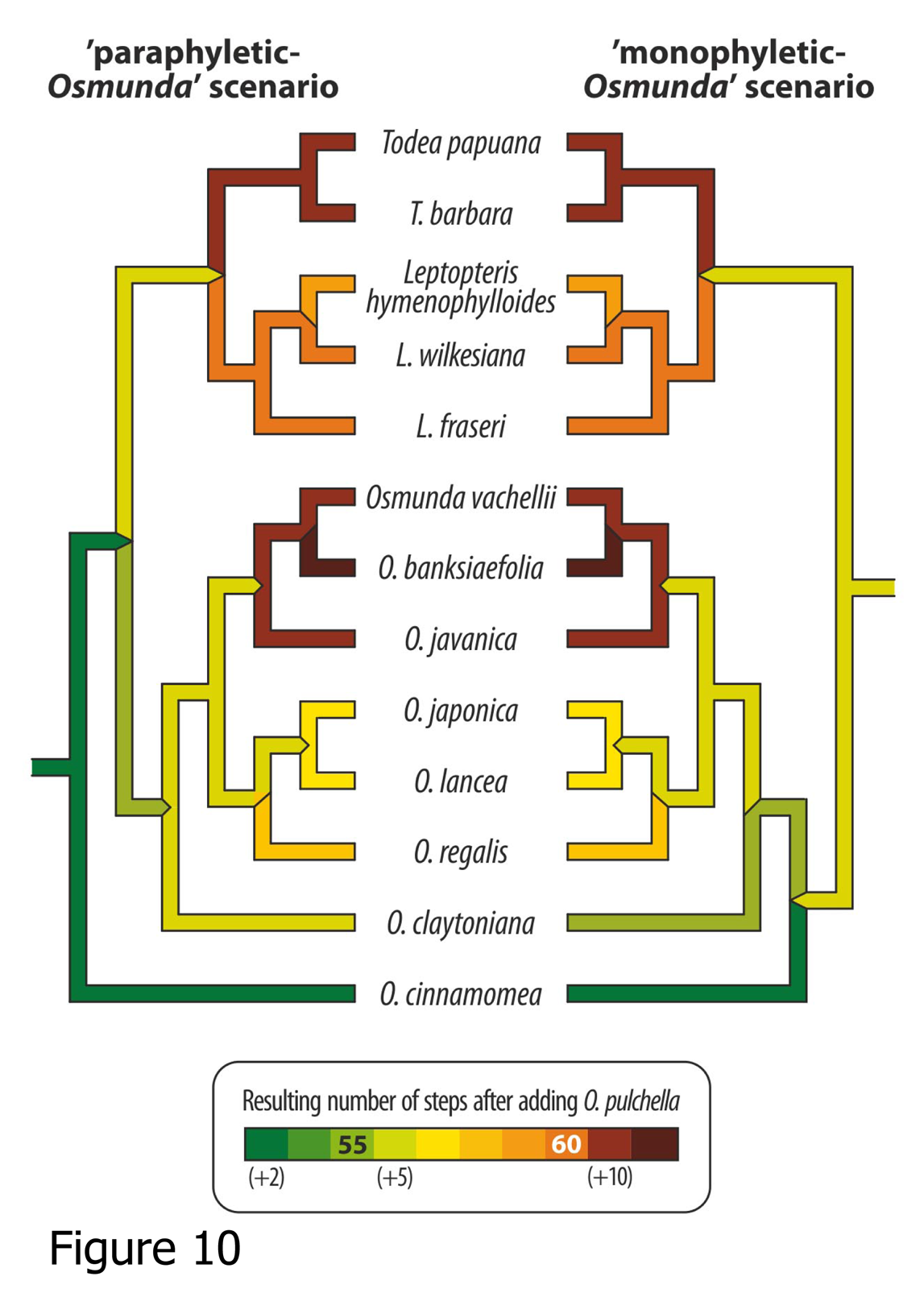
Diagram illustrating the most parsimonious phylogenetic placement of *Osmunda pulchella* within the molecular-based topology under both ingroup rooting scenarios. Left, outgroup-inferred coding gene-based root (Fig. 9; Yatabe et al., 1999; Metzgar et al., 2008). Right, alternative rooting (this study).

## DISCUSSION

In the following sections, we (1) place the new species in the broader context of the fossil record of Osmundaceae; (2) explain the rationale for the assignment of this and other fossil species to an (initially) extant genus; (3) examine the systematic relationships between *Osmunda pulchella* and other fossil and extant species of modern Osmundaceae; (4) consequently, provide a critical re-evaluation of the evidence for generic separation of *Osmundastrum* and the paraphyly of *Osmunda s.l.*; and (5) discuss the impact of the phylogenetic placement of *O. pulchella* on the systematic classification of modern Osmundaceae.

### Osmundaceae in the regional fossil flora

*Osmunda pulchella* sp. nov. is among the earliest fossil *Osmunda* rhizomes known so far, and the first such find from the Mesozoic of Europe. Whole plants are rarely fossilized, so identification of fossils depends on recognizing diagnostic characters in various dispersed organs. Moreover, some isolated organs can only be identified to taxa under special preservational states (e.g., where anatomical details are retained). Fossil evidence for Osmundaceae occurs in three main forms: (1) permineralized axes with vascular, cortical and petiolar anatomy characteristic of the family; (2) compressions and impressions of foliage (either fertile or sterile); (3) dispersed spores with sculptural characters typical of fertile macrofossil or extant representatives of the family.

Permineralized osmundaceous axes have a long-ranging and geographically broad fossil record extending back to at least the Permian of both hemispheres (Gould, 1970; Tian *et al.*, 2008; Wang *et al.*, 2014). These fossils are highly informative of the anatomical evolution of the group since they preserve the three-dimensional architecture of axial tissues and the surrounding sheath of petioles (Miller, 1971). They provide further information on osmundacean ecology, since the excavations or coprolites of various invertebrates are commonly preserved within the cortical tissues or petiole sheath (Tidwell & Clifford, 1995). However, the occurrences of permineralized axes are generally restricted to sedimentary rocks with a high proportion of volcanogenic components. Free silica and, in some cases, carbonate ions are liberated in particularly high concentrations from the breakdown of glass and unstable calc-silicate minerals, especially in sediments derived from mafic to intermediate volcanic terrains (Jefferson, 1987). These ions preferentially link to free hydrogen bonds of holocellulosic complexes in buried plant matter, entombing the original cell walls in opaline silica, quartz, or calcite. The exceptional circumstances of such preservational conditions mean that permineralized osmundaceous stems have a patchy record [see Tidwell (2002) and Tian *et al.* (2008) for summaries of occurrences]. Although axes are known from both older (Permian: Kidston & Gwynne-Vaughan, 1909) and younger (Cenozoic: Kidston and Gwynne Vaughan, 1911; Chandler, 1965; Kvacek & Manum, 1993) rocks in the region, no osmundaceous rhizomes have thus far been reported from the Mesozoic of Europe.

Compressions and impressions of foliage can only be assigned to Osmundaceae with confidence where details of the sori arrangement or sporangial annulus architecture can be resolved (Tidwell & Ash, 1994). Remains of such fertile fronds are variously assigned to *Osmundopsis* T.M.Harris, *Todites* Seward, *Anomopteris* Brongn., *Cacumen* Cantrill & J.A.Webb*, Cladotheca* T.Halle, and *Osmunda* (see Cantrill & Webb, 1987; Tidwell & Ash, 1994; Balme, 1995; Taylor, Taylor & Krings, 2009) and possibly *Damudopteris* D.D. Pant & P.K. Khare and *Dichotomopteris* Maithy (Maithy, 1974a, b). Morphologically similar sterile fronds are typically assigned to *Cladophlebis* Brongn., although not all forms referred to this fossil genus are necessarily osmundacean. Collectively, the record of fossil osmundacean foliage matches that of the rhizomes, extending from the Permian to Cenozoic and being distributed on all continents (Herbst, 1971; Miller, 1971; Anderson & Anderson, 1985; Hill et al, 1999; Collinson, 2001). Foliage referable to *Todites* or *Cladophlebis* is widespread in the Mesozoic of Europe and is extensively represented in Rhaetian to Early Jurassic strata of southern Sweden (Nathorst 1878; Antevs, 1919; Johansson, 1922; Lundblad, 1950; Pott & McLoughlin, 2011).

Spores attributed to Osmundaceae found *in situ* within fossil sporangia or dispersed within sediments are spherical to triangular and typically bear irregularly arranged grana, bacula or pila of variable form and size. More rarely, the spore surface is scabrate or laevigate. When found dispersed, such spores are most commonly assigned to *Osmundacidites* Couper, although some have been attributed to *Baculatisporites* Pflug & P.W.Thomson, *Cyclobaculisporites* D.C.Bhardwaj, *Todisporites* Couper, *Punctatisporites* A.C.Ibrahim, *Leiotriletes* R.Potonié & Kremp, or *Triquitrites* L.R.Wilson & E.A.Coe (Dettmann, 1963; Balme, 1995). Such spores match the record of osmundaceous foliage and permineralized axes in ranging from the Permian to present, and occurring in considerable abundance during the Mesozoic (Balme, 1995). *Osmundacidites wellmanii* (Couper) Danzé-Corsin & Laveine is one of the dominant spore types recovered from sediments surrounding the fossil rhizome studied herein (Bomfleur *et al.*, *in press*) attesting to strong representation of this family in the flora of the Korsaröd area during the Pliensbachian. Moreover, *Osmundacidites* and *Baculatisporites* species are common elements of palynofloras recovered from the uppermost Triassic to Middle Jurassic strata of Sweden (Tralau, 1968; Lund, 1977; Guy-Ohlson, 1986; Lindström and Erlström, 2006; Larsson, 2009), indicating that the family had an important role in the ecology of the herbaceous stratum of the regional mid-Mesozoic vegetation. Osmundaceae underwent a notable decline in both relative diversity and abundance accompanying the rise of the angiosperms in the Cretaceous (Nagalingum *et al.*, 2002; Coiffard, Gomez & Thevenard, 2007) and this trend appears to have persisted through the Cenozoic resulting in the family’s low representation and, for some genera, relictual distribution today (Collinson, 2001).

### Assignment to Osmunda

Historically, permineralized rhizomes similar to those of extant Osmundaceae have been routinely placed in fossil genera, such as *Osmundites* Unger (e.g. Unger, 1854; Kidston & Gwynne-Vaughan, 1910, 1914; see Chandler, 1965; Miller, 1967, 1971). Based on a comparative study of fossil rhizomes and extant taxa, however, Chandler (1965) concluded that *Osmundites dowkeri* Carruth. from the Paleocene of England can be undoubtedly assigned to *Osmunda* subgenus *Plenasium*. Chandler’s rationale has since served as a precedence for subsequent authors to place other Paleogene, Neogene, and—more recently— also Mesozoic fossils of Osmundaceae in genera originally defined for extant species (e.g. Miller, 1967, 1971, 1982; Phipps *et al.*, 1998; Matsumoto & Nishida, 2003; Vavrek, Stockey & Rothwell, 2006; Jud, Rothwell & Stockey, 2008; Carvalho *et al.*, 2013). Finally, well preserved permineralized rhizomes from the Late Cretaceous of Canada that are strikingly similar to those of modern *Osmunda cinnamomea* have led the authors identify particular modern species in even the Mesozoic fossil record (Serbet & Rothwell, 1999). The new combinations and assignments have since been adopted in all systematic treatments of Osmundaceae (e.g. Tidwell & Ash, 1994; Tian *et al.*, 2008, Wang *et al.*, 2014). Hence, the identification of extant genera and species of Osmundaceae even in the Mesozoic fossil record is a unanimously accepted practice, providing the fossils show sufficient diagnostic detail to warrant affiliation with their extant relatives. Fossils that show either insufficient preservation or have structural features unknown among extant taxa, by contrast, continue to be placed in form genera, e.g. *Osmundacaulis* C.N.Mill., *Palaeosmunda* R.E.Gould, *Millerocaulis* Tidwell, *Ashicaulis* Tidwell, and *Aurealcaulis* Tidwell & L.R.Parker (see Tidwell & Ash, 1994; Tian *et al.*, 2008; Wang *et al.*, 2014). Of these, *Ashicaulis* and *Millerocaulis* contain those species that are most similar to extant *Osmunda* [see Vera (2008) for a critical discussion of the generic status of *Ashicaulis* and *Millerocaulis*].

The calcified osmundaceous rhizome described here contains all anatomical features diagnostic of *Osmunda* (see, e.g. Hewitson, 1962; Miller, 1971): (1) ectophloic-dictyoxylic siphonostele with mostly complete leaf gaps; (2) thin parenchymatic inner cortex and distinctly thicker, homogeneous, fibrous outer cortex; (3) heterogeneous sclerenchyma cylinders in the petiole bases; and (4) sclerenchyma fibres in the stipular wings of the petiole. Since the fossil differs from extant species merely in specific diagnostic characters, we have no hesitation in assigning it to *Osmunda* in accordance with conventional practice (see Chandler, 1965; Miller, 1967, 1971, 1982; Tian *et al.*, 2008).

The same also applies to five of the >25 fossil species that are currently included in *Ashicaulis*: *A. liaoningensis* (Zhang & Zheng, 1991), *A. claytoniites* (Cheng, 2011), *A. plumites* (Tian *et al.*, 2014), and *A. wangii* (Tian *et al.*, in press)—all from the Jurassic of China—and *A. johnstonii* from Tasmania (Tidwell, Munzing & Banks, 1991). The holotype of the latter species was collected from a gravel pit; following Tidwell *et al.* (1991), we consider the age of this specimen to be likely concordant with those of other Mesozoic permineralized fern stems from eastern Tasmania, which have recently been dated as Early Jurassic (Bromfield *et al.*, 2007). Consequently, we propose to transfer these species to *Osmunda*, and introduce a replacement name “*Osmunda chengii*” for a resulting junior homonym based on *Ashicaulis claytoniites* (see Appendix).

### Systematic placement of fossil Osmunda rhizomes among modern Osmundaceae

*Phylogenetic network analysis. —* Relationships among extant species in the distance network based on our morphological matrix are congruent with those of molecular phylogenetic analyses (Yatabe *et al.*, 1999; Metzgar *et al.*, 2008), confirming that the morphological matrix based on rhizome anatomy serves well in resolving systematic relationships among modern Osmundaceae. The only major exception is seen in *O. claytoniana*, which, together with extant species of subgenus *Osmunda sensu* Yatabe *et al.* and Paleogene and Neogene fossils, forms a group essentially consistent with subgenus *Osmunda sensu* Miller.

The newly identified Jurassic records of *Osmunda*, including *O. pulchella*, form a wide-spreading box that bridges the gap between the relatively derived *Osmundastrum* and the less derived *Osmunda* subgenus *Osmunda sensu* Miller (Fig. 7). Their long terminal branches are due to unique trait combinations intermediate between their more derived fossil and extant relatives. Collectively, the ‘Jurassic *Osmunda* species’ likely represent ancestral forms of *Osmunda s.l.*, some being more similar to *O. cinnamomea* (*O. pulchella*) and others to subgenus *Osmunda sensu* Miller (e.g. *O. wangii*).

The placement of the other fossil taxa is overall in accordance with the basic assumption that they should be less derived—and thus placed closer to the centre of the network—than their extant relatives. However, there is one major exception: *O. dowkeri* from the Paleogene is the furthest-diverging (i.e. most derived) of all fossil *and* extant species in the *Plenasium* group. This relates to its unusually complex stele organization, which contains by far the largest number of xylem segments of all species analysed (exceeding 30, compared to less than 12 in all other *Plenasium* and less than 20 in most other *Osmunda*.

Notably, a subdivision into two putatively monophyletic subgenera *Osmunda sensu* Yatabe *et al.* and *Claytosmunda* generates two taxa without discriminating anatomical and morphological features (potential aut- or synapomorphies according to Hennig, 1950). Miller’s paraphyletic subgenus *Osmunda* accommodates the fossil taxa, whereas the concept of Yatabe *et al.* (1999, 2005) precludes infrageneric classification of most fossil species (Fig. 7).

*Compatibility with vegetative morphology*. — The systematic relationships revealed from our analysis of anatomical characters of the rhizomes reflect the distribution of gross morphological and fertile features within Osmundaceae very well. The isolated position and tight clustering of subgenus *Plenasium*, for instance, finds support through morphological data in the form of its invariant, unique frond morphology: unlike any other modern Osmundaceae, all extant *Plenasium* species are characterized by having invariably simple-pinnate and hemi-dimorphic fronds. Also the rather wide dispersion of the (paraphyletic) subgenus *Osmunda* Miller is congruent with the variable frond morphology and dimorphism in this group, ranging from pinnate-pinnatifid [e.g. *O. claytoniana* (similar to *O. cinnamomea*)] to fully bipinnate and from fully to variably hemi-dimorphic.

The only major topology where anatomical data alone probably fail to generate a realistic divergence distance occurs in the branch including *Todea* and *Leptopteris*. These genera, with their rhizome anatomy being overall similar to those of *Osmunda* and especially *Osmundastrum* (Hewitson, 1962; but see Fig. 7), have unique vegetative and fertile characters (e.g. isomorphic fronds; tripinnate fronds, arborescent habit, and lack of stomata in *Leptopteris*) that differentiate them very clearly from *Osmunda s.l*.

*Integrating fossil species into the molecular backbone topology. —* The results of the EPA overall provide good support regarding the relationships between fossil and extant taxa (compare Figs 7 and 8). However, notable “position swaps” occur between the placements obtained from different weighting methods of several taxa, including *Osmunda pulchella*. This incongruence is due to intermediate character combinations inherent to ancestral taxa, which we interpret to result in “least conflicting” placements at varying root positions; the EPA is designed to optimize the position of a query taxon within a pre-defined backbone topology. Since *O. pulchella,* and other fossil taxa, show character combinations of genetically distant taxa, the model-based weights in particular will down-weigh the relevant characters. Maximum parsimony has a much more naïve approach in this respect, which may be beneficial for a plausible placement of the fossils. Nevertheless, the fact that this down-weighting results in a placement close to the roots, but not in the tips of sub-trees, indicates that the remaining character suite is plesiomorphic in general, thus supporting the interpretation of fossil taxa such as *O. pulchella* as ancestors of extant clades and possibly species (Figs 7 and 8).

*Summary. —* Altogether, the results detailed above lead us to the following conclusions about the systematic and phylogenetic placements of fossil species among modern Osmundaceae:

1. The Jurassic *Osmunda pulchella* is an ancestral member of *Osmunda s.l.* combining diagnostic features both of *Osmunda s. str.* and of *Osmundastrum*.
2. Other species reported from the Jurassic, together with *O. pluma* (Paleogene) and *O. wehrii* (Neogene), are representatives of the (paraphyletic) subgenus *Osmunda sensu* Miller, including potential ancestors of extant species of subgenus *Osmunda* and *Claytosmunda*.
3. *Osmunda oregonensis* (Paleogene) is closely allied with subgenus *Osmunda sensu* Yatabe *et al.* (see Miller, 1971).
4. *Osmunda arnoldii* and *O. dowkeri* belong to subgenus *Plenasium* and are closely similar to *O. banksiaeafolia*; the highly derived *O. dowkeri* represents the highest degree of specialization in the subgenus, which is supposed to have reached its heyday in distribution and diversity during the Paleogene (Miller, 1971).
5. All fossil *Osmundastrum* can be unambiguously identified as such, despite the wide stratigraphic age-span (Cretaceous, Paleogene, and Neogene) and ‘trans-Pacific’ geographic distribution; it is interesting to note, however, that the rhizomes of *O. cinnamomea* show a far greater disparity in anatomical characters than all other subgenera and even genera of modern Osmundaceae, indicating the existence of probably more than just one *Osmundastrum* species in the past (Fig. 7).
6. *Osmunda iliaensis* and *O. shimokawaensis* are most likely representatives of that species complex of subgenus *Osmunda* that is today restricted to East Asia (i.e. *O. lancea* and *O. japonica*); *O. shimokawaensis* may be ancestral to *O. japonica* and *O. lancea*.
7. the Early Cretaceous *Todea tidwellii* may be as related to modern *Leptopteris* as it is to *Todea*.

### Re-evaluation of the Generic Status of Osmundastrum

The intermediate character combination and the resulting systematic placement of *Osmunda pulchella* and other Jurassic species between *Osmundastrum* and subgenus *Osmunda* Miller as detailed above is incompatible with the current treatment of *Osmundastrum* as a separate genus. In the following section we, therefore, provide a detailed re-evaluation of the sum of evidence that has been used to invoke generic separation of *Osmundastrum*. We begin with what is perhaps considered the most novel and reliable body of evidence—molecular data— and continue with additional evidence from morphological, anatomical, and hybridization studies.

*Molecular data (Fig. 9)*. — The comprehensive multi-locus phylogeny of Metzgar *et al.* (2008) has recently been interpreted to fully support a separate generic status of *Osmundastrum* as suggested by Yatabe *et al.* (1999). Indeed, inter-generic and inter-subgeneric relationships based on the molecular matrix used by Metzgar *et al.* (2008; reproduced here in Fig. 9) receive nearly unambiguous support from the concatenated gene matrix.

However, our root-stability analysis revealed that the inferred paraphyletic status of *Osmunda s.l.* is not unambiguously supported by all gene regions (Fig. 9). Although receiving strong support from the two coding regions (*rbc*L-gene, *atp*A-gene), the molecular data matrix of Metzgar *et al.* (2008) also yields a strong conflicting signal from three relatively conserved spacer sequences (i.e. *trn*L-*trn*F, *atp*B-*rbc*L, and *rbc*L-*acc*D) that indicate an alternative root placement between *Leptopteris*–*Todea* and the remaining *Osmunda s.l. —* offering an equally valid interpretation that would resolve *Osmunda s.l.* as monophyletic.

The root-placement problem may be partly due to the incomprehensive selection of outgroup taxa, which is limited to four samples of leptosporangiate ferns in the matrix of Metzgar *et al.* (2008): *Matonia pectinata* R.Br. (Matoniaceae), *Dipteris conjugata* Reinw. (Dipteridaceae) and *Gleicheniella pectinata* (Willd.) Ching and *Diplopterygium bancroftii* (Hook.) A.R.Sm. (Gleicheniaceae)—all members of Gleicheniales. According to current fern phylogenies, Osmundaceae represent the earliest diverged group in the Polypodiopsida, which include five other extant orders apart from Gleicheniales (see, e.g., Pryer *et al.*, 2004; Smith *et al.*, 2006; Schuettpelz & Pryer, 2007, 2008). In order to obtain a more informative signal, a comprehensive outgroup selection should include taxa from the sister clades of the Polypodiopsida (Marattiopsida and Equisetopsida) and all major lineages within the Polypodiopsida, in particular Hymenophyllales and Schizaeales. Since the Gleicheniales are relatively derived in comparison to the Osmundales, their members may inflict outgroup long-branch attraction with *Osmundastrum* [see Figs S1 (note the long terminal edge bundles) and S2 in ESA].

*Anatomy*. — Rhizomes of extant *O. cinnamomea* exhibit a few peculiar and supposedly unique characters, including (1) the common occurrence of an internal endodermis; (2) the rare occurrence of a dissected, ectophloic to amphiphloic condition of the stele; (3) the protoxylem bundle bifurcating only as the leaf trace enters into the petiole base; (4) the sclerenchyma ring of a petiole base containing one abaxial and two lateral masses of thick-walled fibres; (5) roots arising from the leaf traces usually singly, and only rarely in pairs; and (6) a patch of sclerenchyma adaxial to each leaf trace in the inner cortex (e.g. Hewitson, 1962; Miller, 1971).

The first two characters occur only inconsistently, and are notably absent in Cretaceous to Neogene fossil representatives of *Osmundastrum* (Miller, 1971; Serbet & Rothwell, 1999), indicating that these characters might represent recently acquired traits (Miller, 1971). Moreover, the dissected stele condition, with both endoderms connecting through a leaf gap, occurs only very rarely below incipient rhizome bifurcations, and could only be revealed after thorough investigations of serial sections of over one hundred specimens (Faull, 1901, 1909; Hewitson, 1962). The significance of both characters as diagnostic criteria is thus questionable.

The point of protoxylem bifurcation and the arrangement of patches of thick-walled fibres in the petiole sclerenchyma ring occur consistently, and arguably form appropriate diagnostic criteria for *Osmundastrum*. However, among the remaining *Osmunda s.l.* species, these same two characters are treated as diagnostic features of only specific or subgeneric rank (Miller, 1971); it would thus be inconsistent to weight them more strongly in the delimitation of *Osmundastrum* only.

Roots arising in most cases singly, as opposed to mostly in pairs in the remaining *Osmunda,* is a useful distinguishing character of *Osmundastrum* and *Osmunda pulchella*, although this feature is only inconsistent and may be difficult to observe (Hewitson, 1962; Miller, 1971). The occurrence of sclerenchyma patches adaxial to the leaf traces in the inner stem cortex is the only invariant and unique distinguishing character of *Osmundastrum* that we consider might warrant separation above species level. Apart from *Osmundastrum*, this feature is found only in *Todea* and not in its sister genus *Leptopteris* (Miller, 1971).

*Morphology*. — Morphological features that are commonly cited as diagnostic of *Osmundastrum* include (1) the usually complete frond dimorphism; (2) pinnate-pinnatifid frond architecture; and (3) abaxial hair cover on pinna rachides (Metzgar *et al.*, 2008).

The use of the type of frond architecture and dimorphism as a strict diagnostic character has been shown to be problematic (e.g. Hewitson, 1962). Pinnate frond architecture with deeply pinnatifid segments occurs in both *O.* (*Osmundastrum*) *cinnamomea* and *O.* (*Claytosmunda*) *claytoniana*. In addition, some common varieties and growth forms of *O. cinnamomea* produce only hemi-dimorphic fronds (e.g. Torrey, 1840; Britton, 1890; Kittredge, 1925; Steeves, 1959; Werth, Haskins & Hulburt, 1985), with some having apical fertile portions resembling those of *O. regalis* (see, e.g. Hollick, 1882; Murrill, 1925; Werth *et al.*, 1985) and others having intermittent fertile portions like those of *O. claytoniana* (see, e.g. Day, 1886; Werth *et al.*, 1985). Moreover, completely dimorphic fronds occur also predominantly in *O. lancea*, regularly in *O. japonica*, and sporadically in *O. regalis* (Hooker & Baker, 1883; Chrysler, 1926; see Hewitson, 1962). Notably, such ranges of variation are encountered only in the species complex including *Osmundastrum* and *Osmunda* subgenus *Osmunda* Miller (= subgenera *Claytosmunda* and *Osmunda* Yatabe *et al.*).

Finally, fronds of all *Osmunda s.l.* species emerge with a more-or-less dense abaxial indumentum and differ merely in the duration to which the trichome cover is retained in the course of frond maturation (Hewitson, 1962). In fully mature fronds of all species considered, most of the hair cover is ultimately lost, with *O. cinnamomea* [especially *O. cinnamomea* var. *glandulosa* Waters (see Waters, 1902; McAvoy, 2011)] merely tending to retain greater amounts of hairs than *O. claytoniana*, and those in turn more than other species (Hewitson, 1962). In summary, we follow Hewitson (1962) and consider none of these morphological features to provide consistent and reliable diagnostic characters for separating *Osmundastrum* from subgenus *Osmunda* Miller.

*Hybridization*. — Metzgar *et al.* (2008, p. 34) suggested that the existence of hybrids can be used to decide about the elevation of subgenera to generic ranks. A range of natural hybrids, intra- and inter-subgeneric, are known to occur in *Osmunda s. str.*: *O.* × *ruggii* R.M.Tryon in eastern North America (*O. regalis* × *O. claytoniana*; Tryon, 1940; Wagner *et al.*, 1978), *O.* × *mildei* C.Chr. in southern China (*O. japonica* × *O. vachellii* Hook.; Zhang *et al.*, 2008; Kato *et al.*, 2009), *O*. × *hybrida* Tsutsumi, S.Matsumoto, Y.Yatabe, Y.Hiray. & M.Kato in Southeast Asia (*O. regalis* × *O. japonica*; Tsutsumi *et al.*, 2011), and *O*. × *intermedia* (Honda) Sugim. (*O. japonica* × *O. lancea*) and *O.* × *nipponica* Makino (*O. japonica* × ?*O. claytoniana*) in Japan (Kato, 2009; Yatabe *et al.*, 2009; Tsutsumi *et al.*, 2012). The seeming absence of naturally occurring hybrids involving *Osmundastrum* has been interpreted to result from its particularly isolated position within *Osmunda s.l.* (Miller, 1967, 1971). However, Klekowski (1971) conducted artificial breeding experiments and readily succeeded in producing viable hybrid sporophytes from *O. cinnamomea* × *O. claytoniana* and *O. cinnamomea* × *O. regalis*, with equal or even higher yields (1 out of 8 and 2 out of 9, respectively) compared to *O. claytoniana* × *O. regalis* (1 out of 8). In addition, some authors suspect that there may also be natural hybrids between *O. cinnamomea* and *Osmunda s. str.* (e.g. Sugimoto, 1979). So far, there is no record about *in* or *ex situ* hybridisation between *Leptopteris-Todea* and *Osmunda s.l*.

*Summary*. — We find that neither molecular, anatomical, morphological, nor hybridization studies have yet succeeded in providing unequivocal evidence that would warrant separate generic status of *O. cinnamomea*. We argue that the sum of evidence for extant taxa detailed above rather allows for two equally valid interpretations: the ‘paraphyletic-*Osmunda* scenario’ (Yatabe *et al.*, 1999; Metzgar *et al.*, 2008) and an alternative ‘monophyletic-*Osmunda* scenario’ (e.g. Miller, 1971).

### The impact of Osmunda pulchella on the classification of modern Osmundaceae (Figs 10, 11)

The phylogenetic placement of *Osmunda pulchella* is critical to the systematic classification of modern Osmundaceae. In the specified topology of the ‘paraphyletic *Osmunda* scenario’, most parsimonious placement of *O. pulchella* is at the base of the tree, at the root of either *Osmundastrum* or of the remaining *Todea*-*Leptopteris-Osmunda s. str.* clade (Fig. 10). If this phylogenetic scenario is followed, and if only monophyletic groups are considered valid taxonomic units (see, e.g. Hörandl, 2007 and Hörandl & Stuessy, 2010, for critical discussion), then it follows that all modern Osmundaceae need be included in one genus *Osmunda*, with *Plenasium*, *Osmunda/Claytosmunda*, *Osmundastrum*, *Todea*, and *Leptopteris* being infrageneric taxa (Fig. 11).

**FIG. 11.**
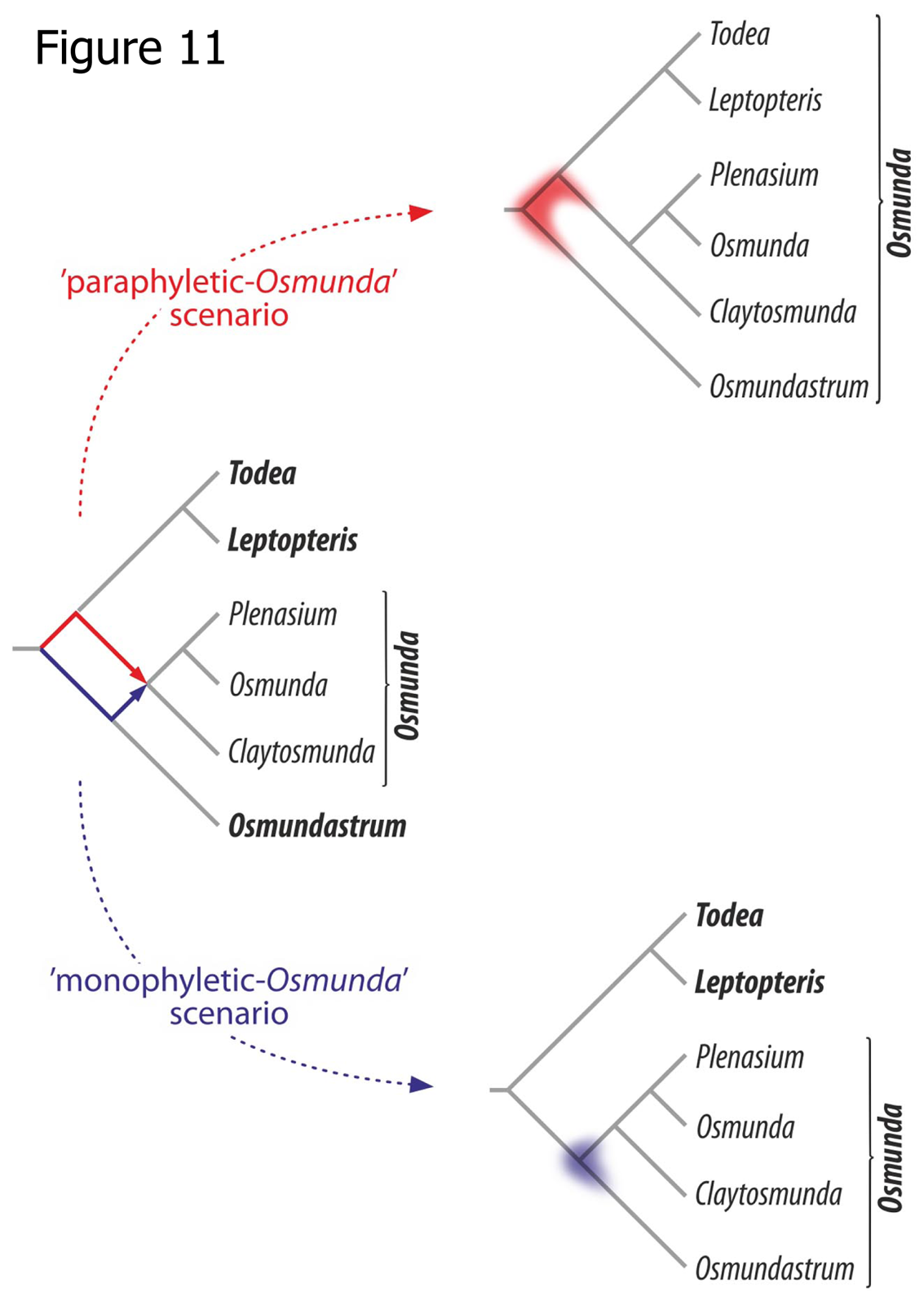
Diagram illustrating the critical significance of the placement of *Osmunda pulchella* (colour shading) for a strictly cladistic-systematic classification of modern Osmundaceae regarding the two alternative rooting schemes (Figs 9, 10). Genus names in bold; infrageneric taxon names in regular font.

If, however, the specified topology of the ‘monophyletic *Osmunda* scenario’ is followed, in which most parsimonious placement of *O. pulchella* is as sister to *O. cinnamomea* at the base of an *Osmundastrum-Osmunda s. str.* clade (Fig. 10), then all fossil and extant species of modern Osmundaceae can be resolved in three mutually monophyletic genera: *Todea*, *Leptopteris*, and *Osmunda* including the subgenera *Plenasium*, *Osmunda*, *Claytosmunda*, and *Osmundastrum* (Fig. 11).

In our opinion, this latter option integrates the seemingly conflicting evidence from studies of the morphology, anatomy, molecular data, and fossil record of Osmundaceae in a much more realistic and elegant way, and—beyond that—offers a more practical taxonomic solution. We, therefore, argue that *Osmunda pulchella* described here exposes the recently established paraphyly of *Osmunda s.l.* as a result of a sampling or reconstruction artefact in the molecular matrix employed.

## SUMMARY AND CONCLUSIONS

i. *Osmunda pulchella* sp. nov. from the Early Jurassic of Sweden is among the earliest unequivocal records of fossil *Osmunda* rhizomes.
ii. Analogous to our treatment of *O. pulchella*, five additional species from the Jurassic of China and Australia, currently assigned to the form-genus *Ashicaulis*, show all diagnostic features of *Osmunda s.l.* and are accordingly transferred to that genus: *O. chengii* (based on *A. claytoniites*), *O. johnstonii*, *O. liaoningensis*, *O. plumites*, and *O. wangii*.
iii. Intermediate anatomical character suites of Jurassic *Osmunda* species support re-inclusion of the recently separated, monospecific *Osmundastrum* into *Osmunda*.
iv. The sum of morphological, anatomical, molecular, and fossil evidence supports modern *Osmunda* (including *Osmundastrum*) and *Todea-Leptopteris* being mutually monophyletic.
v. The recently established rooting of Osmundaceae and the resulting paraphyly of *Osmunda s.l.*, based solely on molecular data, likely results from a sampling or reconstruction artefact.

## SUPPLEMENTARY INFORMATION

Supplementary Information to this article is available online on the journals homepage and consists of a document file (File_S1_DetailedMethods.doc) containing an annotated list of the characters in the morphological matrix, the character coding of included taxa, and a detailed description of methods employed in the phylogenetic reconstructions; Figure S1 (Fig_S1_MolecDistNetwork.pdf), showing systematic relationships among extant Osmundaceae in the form of a neighbour-net inferred from uncorrected pairwise distances based on the concatenated data set of Metzgar *et al.* (2008); Figures S2 and S3 (Fig_S2_HeatMapMolecularMatrix.pdf, Fig_S3_HeatMapMorphMatrix.pdf), containing heat maps showing general similarity patterns between taxa in the molecular and the morphological data matrices, respectively.

In addition, an electronic supplementary data archive (ESA) is available online at www.palaeogrimm.org/data/Bfr14_ESA.zip, consisting of the following: File S1, Figures S1– S3; File S2 (File_S2_MorphologicalFeatures.xlsx), containing a spreadsheet with a compilation of morphological data used in the matrix; and three folders (labelled “ClustQuantChars”, “Inferences”, and “Matrices”) containing all original data files, including the employed matrices in NEXUS format [please refer to the accompanying index document (GuideToFiles.txt) for a detailed description].

## ACKNOWLEDGEMENTS

We thank Wang Yongdong (Nanjing) and Tian Ning (Shenyang) for discussion and assistance in obtaining literature; Alexandros Stamatakis (Heidelberg) for discussing methodological approaches; Birgitta Bremer and Gunvor Larsson (Stockholm) for providing live material of *Osmunda* for comparison; and Else Marie Friis and Stefan Bengtson (Stockholm), Christopher H. Haufler (Lawrence, KS), and Troy E. Wood (Flagstaff, AZ) for discussion.

## APPENDIX

### Taxonomic treatment of selected species previously assigned to Ashicaulis

For reasons detailed in the discussion and by analogy with the systematic placement of *Osmunda pulchella*, we propose to transfer five species that are currently accommodated in fossil form-genera (*Millerocaulis* and *Ashicaulis*) to *Osmunda*.

Order Osmundales Link

Family Osmundaceae Berchtold & Presl

Genus *Osmunda* L.

*Osmunda chengii* Bomfleur, G. Grimm & McLoughlin, nom. nov.

*Basionym. — Ashicaulis claytoniites* Y.M. Cheng (in Review of Palaeobotany and Palynology 165: 98. 2011)

*Synonym (replaced)*. — *Osmunda claytoniites* (Y.M. Cheng) Bomfleur, G. Grimm & McLoughlin, comb. nov.

[*Senior homonym*. — *Osmunda claytoniites* Carlie J. Phipps, T.N. Taylor, Ed.L. Taylor, Cúneo, L.D. Boucher & X. Yao (in American Journal of Botany 85: 889. 1998)]

*Remarks:* The resulting new combination is a junior homonym of *Osmunda claytoniites* from the Triassic of Antarctica (Phipps *et al.*, 1998). In accordance with Articles *6.10*, *6.11*, and *41* of the International Code of Nomenclature for algae, fungi, and plants (Melbourne Code, 2011), we propose the replacement name *Osmunda chengii*. The specific epithet is chosen in honour of Cheng Ye-Ming (Beijing, China), author of the original species name.

*Osmunda johnstonii* (Tidwell, Munzing & M.R. Banks) Bomfleur, G. Grimm & McLoughlin, comb. nov.

*Basionym*. — *Millerocaulis johnstonii* Tidwell, Munzing & M.R. Banks (in Palaeontographica B 223: 94. 1991); see also Vera (2008)

*Synonym*. — *Ashicaulis johnstonii* (Tidwell, Munzing & M.R. Banks) Tidwell (in SIDA 16: 256. 1994)

*Osmunda liaoningensis* (Wu Zhang & Shao-Lin Zheng) Bomfleur, G. Grimm & McLoughlin, comb. nov.

*Basionym*. — *Millerocaulis liaoningensis* Wu Zhang & Shao-Lin Zheng (in Acta Palaeontologica Sinica 30: 717. 1991); see also Vera (2008).

*Synonym*. — *Ashicaulis liaoningensis* (Wu Zhang & Shao-Lin Zheng) Tidwell (in SIDA 16: 256. 1994) (authorities for the basionym and the new combination are cited as “Wu & Shao Lin”).

*Osmunda plumites* (N. Tian & Y.D. Wang) Bomfleur, G. Grimm & McLoughlin, comb. nov. *Basionym*. — *Ashicaulis plumites* N. Tian & Y.D. Wang (in Journal of Plant Research 127: 210. 2014)

*Osmunda wangii* (N. Tian & Y.D. Wang) Bomfleur, G. Grimm & McLoughlin, comb. nov. *Basionym*. — *Ashicaulis wangii* N. Tian & Y.D. Wang (in Science China: Earth Sciences xxx: xxx. in press)

